# Senescent cell death as an aging resistance mechanism in naked mole-rat

**DOI:** 10.1101/2020.07.02.155903

**Authors:** Yoshimi Kawamura, Kaori Oka, Mayuko Takamori, Yuki Sugiura, Yuki Oiwa, Shusuke Fujioka, Sayuri Homma, Shingo Miyawaki, Minoru Narita, Takaichi Fukuda, Makoto Suematsu, Hidemasa Bono, Hideyuki Okano, Kyoko Miura

## Abstract

Naked mole-rats (NMRs) are the longest-lived rodents, showing minimal aging phenotypes. An unsolved paradox is that NMRs exhibit low intracellular anti-oxidant defence despite minimal aging. Here, we explained a link between these “contradicting” features by a phenomenon termed “senescent cell death (SCD)”—Senescence induced cell death in NMR cells due to their inherent vulnerability to reactive oxygen species and unique metabolic system. In NMR skin, we observed few senescent cells during aging or after ultraviolet irradiation, suggesting suppression of senescent cell accumulation in NMR tissue. We discovered that senescent NMR-fibroblasts induce SCD through retinoblastoma protein activation accompanied by autophagy dysregulation, increased oxidative damage and accelerated H_2_O_2_-releasing metabolic pathways. During senescence, NMR cells showed resistance to metabolic remodelling unlike mice. Our findings provide mechanistic insights into how extraordinary aging resistance is accomplished in NMR. This will contribute to the development of senolytic drugs to regulate age-related diseases.

---

Naked mole-rats (*Heterocephalus glaber*, NMRs) are African eusocial mammals which form a unique social structure similar to ants or bees^1^. NMRs live in underground tunnels with few air vents, a hypoxic environment in which oxygen concentration sometimes drops to 6–7%. NMRs exhibit extraordinary longevity with a maximum lifespan of 32 years, although NMR body mass is similar to that of the mouse^2^. Moreover, NMRs show a unique senescence phenotype^2^. NMR defies the Gompertzian laws of mortality; i.e. NMR mortality rate does not increase during lifetime, and body functions including fecundity are maintained throughout the lifetime^3^. Moreover, NMRs have unusual resistance to several age-related diseases such as metabolic disorders, neurodegenerative disease, and cancer^4–6^. Recently, several mechanisms that enable NMR’s longevity, delayed aging, and cancer-resistance phenotypes are proposed^7–12^.

Generally, the “free radical theory of aging”, later modified to “mitochondrial free radical theory”, is the well-known theory of aging mechanism^13^. Intracellular reactive oxygen species (ROS), deriving especially from mitochondria, damages macromolecules such as lipids, DNA, and proteins, and the accumulated damages in tissues are assumed to contribute aging process^14^. Indeed, the mitochondrial ROS production rate is negatively correlated with the maximal lifespan of animal species^15,16^.

However, previous insights on responses of long-lived NMRs to ROS are puzzling: 1) Several reports suggested that NMRs have stronger anti-oxidant mechanism(s). In NMR tissues, mitochondrial hydrogen peroxide (H_2_O_2_) generation rate was similar to mice but the mitochondrial ROS consuming capacity was higher in NMRs^17^. Similarly, the vascular O_2^-^_· and H_2_O_2_ production in NMR blood vessels was comparable to some rodent species, but cultured NMR vessels showed resistance to H_2_O_2_-induced cell death^18^. In addition, the nuclear factor erythroid 2-related factor 2 (NRF2) signalling pathway is activated in NMRs, which upregulates various cytoprotective genes including ROS-responding genes^11^. 2) However, many other reports suggested that NMRs exhibit low anti-oxidant defence. The activity of H_2_O_2_ removal enzyme (glutathione peroxidase, gpx) in NMR liver was 70 times lower than mice liver^19^. In addition, NMR fibroblasts are highly susceptible to H_2_O_2_ treatment but resistant to paraquat treatment as compared to mouse fibroblasts^20,21^. From young ages, NMR suffers greater oxidative damages in tissue DNA, protein, and lipids than mice^10,22^. Nevertheless, the level of oxidative damage does not increase further and remains constant for more than 20 years^10^. Thus, at least in part, NMR exhibits low intracellular anti-oxidant defence despite their delayed aging. It has been considered that the delayed aging of NMRs, although suffering high oxidative damage, may be due to NRF2 signalling activation and high protein stability^10,23^. On the other hand, there are currently no reports that directly link the NMR’s intracellular vulnerability to ROS, and NMR’s longevity or delayed-aging phenotype. These complex but interesting observations raise a possibility that NMR may have developed a unique system to remove damaged cellular components or the cells that suffered the oxidative damage during aging.

In mammalian cells, one of the typical “damaged” cellular status along with elevated oxidative damage is cellular senescence. Cellular senescence is an irreversible cell proliferation arrest induced in response to stresses such as DNA damage, oncogene activation, and telomere shortening^24^. Cellular senescence contributes to avoid cancer formation by stopping proliferation of damaged cells^24^. In addition, cellular senescence has important roles in tissue homeostasis, embryonic development and wound healing^25–27^. On the other hand, accumulation of senescent cells promotes age-related physiological deterioration and disorders, by secreting a bioactive “secretome” called senescence-associated secretory phenotype (SASP). SASP includes pro-inflammatory cytokines, chemokines, growth modulators, angiogenic factors, and matrix metalloproteinases^24,28^. Furthermore, in senescent cells, cell-intrinsic stressors increase^29,30^. A typical stressor is intracellular ROS. ROS elevation plays a pivotal role in development and maintenance of senescence state through DNA damage and activation of persistent DNA damage response^31,32^. Whereas non-senescent cells would normally go into apoptosis upon increased intracellular stress, senescent cells avoid cell death^33^. Senescent cell accumulation during aging likely plays a causative role in aging and age-related diseases, as clearance of senescent cells delays aging and age-related disorders^34,35^.

Several reports demonstrated that NMR cells had the potential to become senescent. Although NMRs do not show replicative senescence^36^, Zhao *et al*. recently showed that cellular senescence was observed in NMR during the developmental process or when received stresses such as DNA damage or oncogene activation^37^. We have previously shown that *ARF* (a tumour suppressor gene) suppression in stressed NMR fibroblasts induces cellular senescence-like state, termed *ARF*-suppression induced senescence (ASIS)^12^. These observations indicate that NMRs can undergo cellular senescence in several situations. However, it is still unclear whether senescent cells accumulate in the NMR body during aging.

The objective of our study is to investigate aging mechanism in longest-lived rodents NMRs by establishing a link between low intracellular anti-oxidant defence despite minimal aging phenotype. Our research examined the connection between delayed aging and vulnerability to ROS focusing on the response to cellular senescence in NMR. In this study we attempted to shed light on the novel mechanism of extraordinary aging resistance in NMRs.

## Results

### Accumulation of senescent cells is highly suppressed in NMR skin tissues during aging and after UV irradiation

To determine whether senescent cells accumulate in NMR skin tissues during aging, we collected skin biopsy specimens from 13–15 year-old NMRs (middle-aged, the oldest available animals) and one year-old NMRs (young) (Fig. 1a). As controls, we also collected skin biopsies from one year-old mice (middle-aged) and four week-old young mice (young). In middle-aged NMR skin (13–15 year-old), we found almost no cells positive for senescence-associated beta-galactosidase (SA-β-Gal) activity, which is associated with cellular senescence. On the other hand, middle-aged mouse skins (one year-old) showed a significant increase in SA-β-Gal-positive cells (Fig. 1b, c). The brown dots in dermis of NMR skin are melanin pigments, a common feature in relatively young NMRs (Fig. 1b, e). Moreover, quantitative reverse transcription-polymerase chain reaction (qRT-PCR) showed that, in middle-aged NMR skins, age-associated increase in expression of senescence-associated marker *INK4a* was much less than in middle-aged mice. The expression level of another senescence-associated marker *p21* was rather decreased with age in NMRs (Fig. 1d). These results indicate that NMRs have resistance to senescent cell accumulation during aging process. On the other hand, SA-β-Gal-positive cells were observed in digits of neonate NMRs, indicating that developmental senescence occurs in NMR tissue as previously described^37^ (Supplementary Fig. 1a). Next, we experimentally induced cellular senescence in skin by UV-B irradiation as previously described (Supplementary Fig. 1b)^38^. Although many SA-β-Gal positive cells appeared in dermis of mouse skin, we did not observe accumulation of SA-β-Gal positive cells in NMR skins after UV-B treatment (Fig. 1e, f). Notably, in contrast to mouse skins, after UV-B irradiation the cells positive for cell death markers (like TUNEL staining or cleaved Caspase-3) were significantly increased in NMR skin dermis (Supplementary Fig. 1c-f). These results demonstrate that the NMR skin tissues hardly accumulate senescent cells during aging or after receiving DNA damage, which led us to ask whether the fate of NMR cells during cellular senescence is different from the fate of cells in other mammalian species.

**Figure 1.**
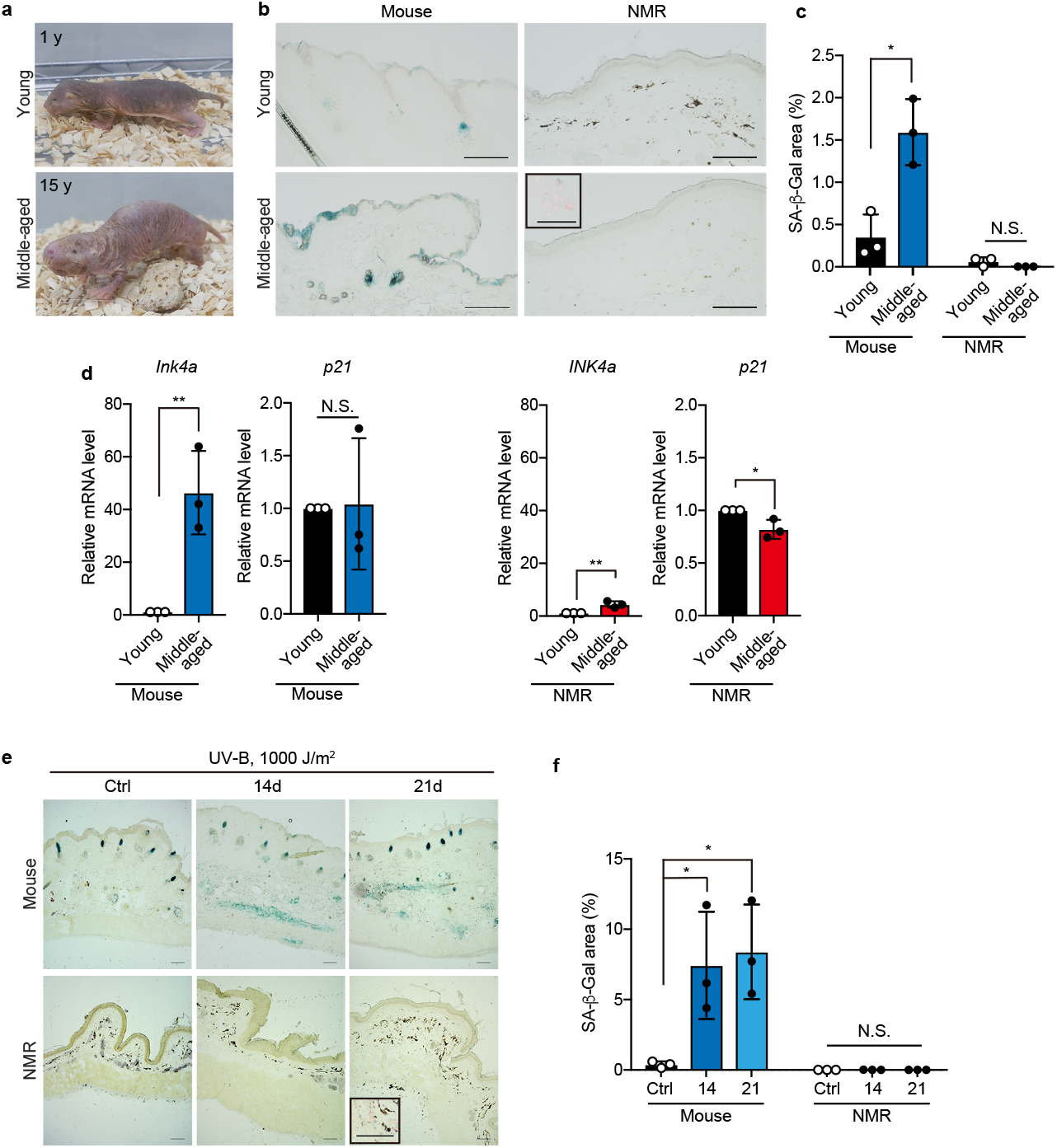
NMR skin hardly accumulates senescent cells during aging or by UV irradiation. **a**, Young NMR (one year-old) or middle-aged NMR (15 year-old). **b**, SA-β-Gal activity in NMR- or mouse-skin. Four week-old young mice (young), one year-old aged mice (middle-aged), one year-old young NMRs (young) and 13-15 year-old aged NMRs (middle-aged) were used. Rectangle in the middle-aged NMR panel indicates SA-β-positive cells and nuclei (red). The brown dots in dermis of NMR skin are melanin pigments. Scale bar, 100 μm. **c**, Quantification of SA-β-Gal-positive area (%). **d**, qRT-PCR analysis of the expression of *INK4a* and *p21* in the skin of each group. **e**, SA-β-Gal activity in skins of NMR (young, one year-old) or mouse (young, 6 weeks-old) after 1000 J/m^2^ of UV-B irradiation. Rectangle in the NMR skin panel at 21 days after irradiation indicates SA-β-positive cells. The brown dots in dermis of NMR skin are melanin pigments. Scale bar, 100 μm. **f**, Quantification of SA-β-Gal-positive area (%) in the skins after UV-B irradiation. * *P* < 0.05; ** *P* < 0.01; N.S. not significant, unpaired *t*-test versus control for **c** and **d**; One-way analysis of variance (ANOVA) followed by Dunnett’s multiple comparison test for **f**. Data are mean ± SD from *n* = 3 animals respectively.

### Senescent NMR fibroblasts gradually activate cell death through RB activation but not through p53

To elucidate the cellular fate of NMR fibroblasts during senescence, we generated primary skin fibroblasts and induced cellular senescence *in vitro*. As cellular senescence induction by damaging DNA also causes acute cell death in proliferative non-senescent cells^33^, we instead performed lentiviral transduction of INK4a into fibroblasts of NMRs or mice to analyse the fate of senescent cells (Supplementary Fig. 2a, b). INK4a is a cyclin-dependent kinase inhibitor, and under normal culture condition, INK4a transduction efficiently induces cellular senescence by inhibiting cyclin-dependent kinases CDK4/6 and activating Retinoblastoma family (RB)^32,39^. 12 days after INK4a transduction, several features of cellular senescence, such as enlarged and flat cell morphology, decrease in BrdU incorporation, increase in SA-β-Gal activity, hypophosphorylation of RB protein, phosphorylation of AKT protein, and increase in g-H2AX and 53BP1 foci number were observed in both NMR and mouse fibroblasts (Fig. 2a–f and Supplementary Fig. 2c). Although NMR’s body temperature is about 32 °C, there was no significant difference in SA-β-Gal activity in NMR fibroblasts between 32 °C and 37 °C (Supplementary Fig. 2d).

**Figure 2.**
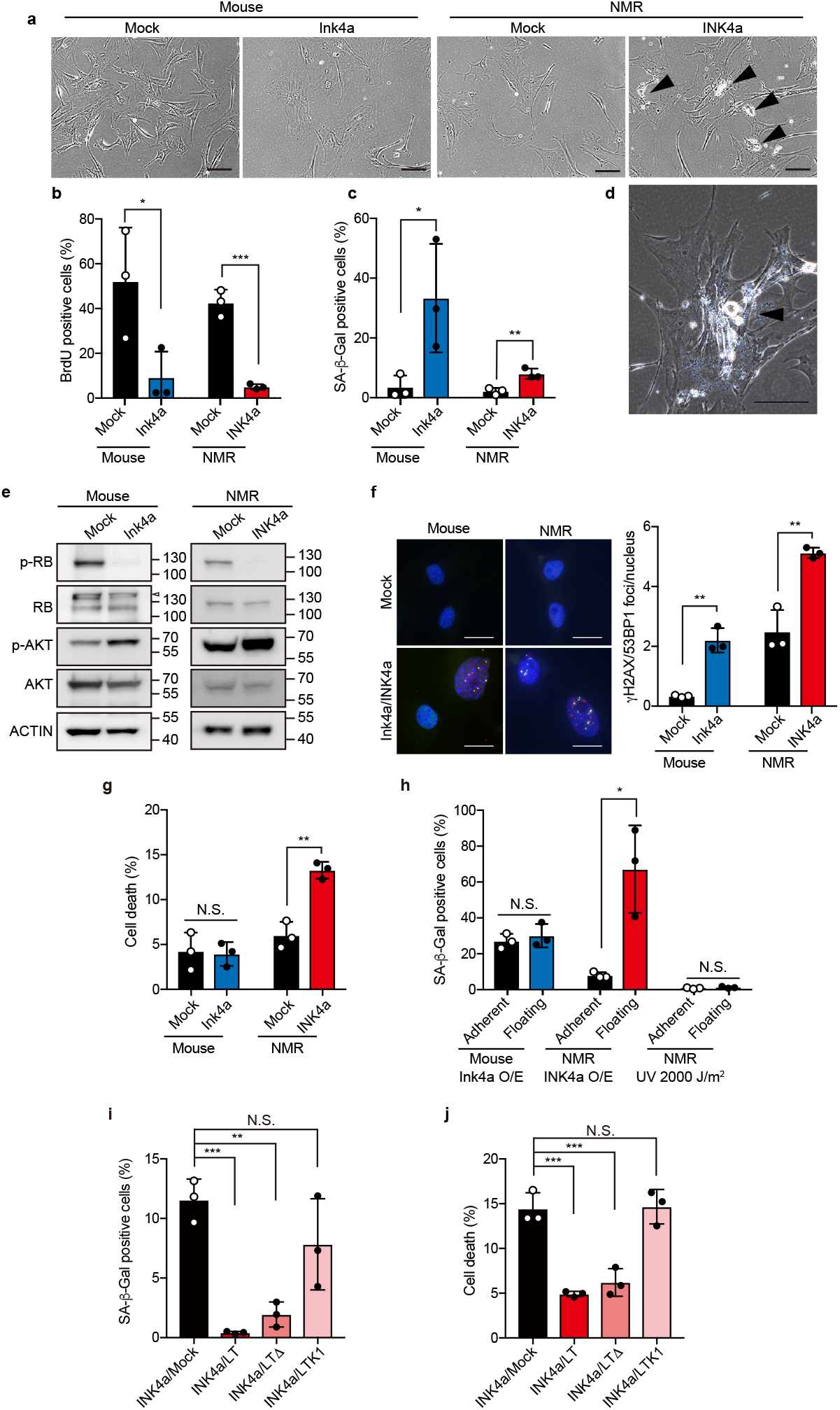
Senescent NMR fibroblasts gradually activate cell death through RB but not p53. **a**, Cell morphology of mouse- or NMR-fibroblasts 12 days after INK4a transduction. Arrow heads indicate dying cells. Scale bar, 100 μm. **b**, Quantification of BrdU-positive cells (%). **c**, Quantification of SA-β-Gal-positive cells (%). **d**, High-magnification image of NMR fibroblasts 20 days after INK4a transduction. Arrow head indicates dying cells. Scale bar, 100 μm. **e**, Western blot of RB and AKT in NMR- or mouse-fibroblasts at 12 days after INK4a transduction (p, phosphospecific antibody). ACTIN was used as a loading control. Arrowhead indicates a non-specific band. **f**, Left, representative images of g-H2AX (red) and 53BP1 (green) staining in mouse- or NMR-skin fibroblasts at 12 days after INK4a transduction. Scale bar, 20 μm. Right, quantification of co-localized g-H2AX and 53BP1 foci. **g**, Quantification of Annexin V-positive cells (%). **h**, Quantification of SA-β-Gal-positive cells in floating dead cell population and adherent living cell population in mouse- or NMR-fibroblast culture. **i**, Quantification of SA-β-Gal-positive cells (%) in NMR-fibroblasts transduced with different forms of SV40 Large T antigen (LT, LTD, and LTK1) and INK4a. **j**, Quantification of Annexin V-positive cells (%). * *P* < 0.05; ** *P* < 0.01; *** *P* < 0.001; Unpaired *t*-test versus control for **b, c, f, g** and **h**. One-way ANOVA followed by Dunnett’s multiple comparison test for i and j. Data are mean ± SD from *n* = 3 biological replicates.

Interestingly, only in NMR fibroblasts 12 days after INK4a transduction, we found a lot of dead floating cells (arrowheads in Fig. 2a). Annexin V/PI staining showed that cell death was significantly increased only in NMR fibroblasts and not in mouse fibroblasts 12 days after INK4a transduction (Fig. 2g and Supplementary Fig. 2e). At day 20, cell death was further enhanced in INK4a-transduced NMR fibroblast culture (Fig. 2d and Supplementary Fig. 2e). Notably, after the INK4a transduction, we observed that SA-β-Gal-positive NMR cells were significantly enriched in the floating, dead cell population but not in the live adherent cell population (Fig. 2h and Supplementary Fig. 2f). By contrast, in mouse fibroblasts, Ink4a transduction did not cause such enrichment of SA-β-Gal-positive cells. As intense UV irradiation led NMR fibroblasts into acute cell death and did not make the cells SA-β-Gal-positive, high SA-β-Gal activity is not a common phenotype of dead NMR cells (Fig. 2h and Supplementary Fig. 2f). The time-course analysis showed that activation of cell death was correlated with upregulation of INK4a expression and SA-β-Gal activity in NMR fibroblasts after INK4a transduction (Supplementary Fig. 2g–i). Notably, NMR fibroblasts derived from lung also showed increase in cell death at 12 days after INK4a transduction (Supplementary Fig. 2j–l).

To evaluate the response of NMR cells to other cellular senescence-inducing stimuli, we treated fibroblasts with a low concentration of mitomycin C (MMC), a DNA-damaging alkylating agent that induces premature senescence at low concentration^40^. 10 days after the MMC treatment, both mouse and NMR fibroblasts upregulated cellular senescence markers, however, only in NMR fibroblasts but not in mouse-fibroblasts, the activation of cell death was observed (Supplementary Fig. 3a–e). Similar to the above mentioned INK4a-transduced NMR fibroblast model, a significant enrichment of SA-β-Gal-positive cells occurred in the floating dead cells but not in adherent live cells in MMC-treated NMR fibroblasts; indicating that mainly senescent cells go into cell death (Supplementary Fig. 3f). Knockdown of INK4a induced a trend towards cell death attenuation in MMC-treated senescent NMR fibroblasts, although not statistically significant (Supplementary Fig. 3g, h). Moreover, NMR fibroblasts transduced with HRasV12, which causes oncogene-induced senescence^41^, also resulted in similar activation of cell death (Supplementary Fig. 3i–m). These results demonstrate that NMR cells show a unique phenotype in which cells gradually go into cell death during senescence. We termed this phenomenon as senescent cell death (SCD).

In mammalian cells, activations of Ink4a-Rb and p53-p21 pathways are important for senescence induction. Also, high level of p53 activation leads to apoptotic genes transcription and ensuing apoptosis^33^. To clarify whether SCD in NMR cells requires activation of RB and/or p53 pathways, we used SV40 Large T antigen (LT) and its derivatives to distinguish activities of RB and p53 pathways. Wild-type LT is a viral oncoprotein that suppresses both p53 and RB (p53-/RB-). LTΔ434-444 (LTD) mutant inactivates only RB (p53+/RB-). LTK1 mutant suppresses only p53 (p53-/RB+)^42^. We transduced each of these proteins together with INK4a into NMR fibroblasts by lentiviral vector (Supplementary Fig. 4a). As a result, wild-type LT (p53-/RB-) and LTD (p53+/RB-) strongly suppressed SA-β-Gal activity and cell death (Fig. 2i, j, Supplementary Fig. 4b). On the other hand, inactivation of only p53 by LTK1 (p53-/RB+) did not affect SCD in INK4a-transduced NMR fibroblasts; while expression of p21, a p53 downstream gene, was reduced (Fig. 2i, j, Supplementary Fig. 4c). When LTD (p53+/RB-) was introduced, the expression of p21 increased (Supplementary Fig. 4c). This may be due to a compensatory increase of p53 activity induced by RB inactivation. INK4a, which caused SCD in NMR cells, is a cyclin-dependent kinase inhibitor (CKI) that activates RB. Thus, we next evaluated whether SCD is caused by other RB-activating CKIs such as p15, p21, p27, and pALT (a hybrid isoform of p15 and INK4a expressed in NMRs)^43^. The transduction of CKIs other than INK4a also resulted in activation of RB, increased SA-β-Gal positive cells, and significant activation of cell death (Supplementary Fig. 4d–h). Collectively, these results demonstrate that activation of RB, but not p53, is required for the induction of SCD in NMR fibroblasts.

A previous report showed that species-specific production of high-molecular-mass hyaluronan increased the expression of INK4a and contributed to early contact inhibition phenotype of NMR fibroblasts^8^. Thus, we next assessed whether high-molecular-mass hyaluronan contributed to SCD in NMR fibroblasts by transduction of INK4a in the presence of *Streptomyces* hyalulonidase (HAase). However, we did not observe any change in SCD (Supplementary Fig. 4i).

### Dysregulation of autophagy and increase of oxidative stress in senescent NMR cells

Generally, senescent cells acquire resistance to apoptosis by activation of anti-apoptotic BCL2 family such as BCL2, BCL-XL, and BCL-W^44^. Because INK4a-transduction in NMR fibroblasts did not upregulate expression of BCL2 family genes (Supplementary Fig. 5a), we first focused on the possibility that inefficient anti-apoptotic protein upregulation in senescent NMR fibroblasts might contribute to SCD. However, transduction of BCL-XL or treatment with caspase inhibitor Z-VAD-FMK were not able to suppress SCD (Supplementary Fig. 5b–d). Thus, the lack in up-regulation of anti-apoptotic proteins is not the major cause of SCD in NMRs.

Next, in order to explore the mechanism of SCD, we performed a comparison of global gene expression between senescent and non-senescent NMR fibroblasts by RNA-sequencing (RNA-seq) (Fig. 3a). Upregulated genes (>1.5-fold) in INK4a-transduced NMR cells compared to mock-transduced NMR cells were analysed to determine the enriched pathways using Metascape^45^. Top-ranked 20 enriched gene ontology (GO) terms and KEGG pathways are shown in Fig. 3b. The gene names in each enriched pathway are summarized in Supplementary Table 1. By INK4a transduction, a significant enrichment of genes related to “SASP”, “aging”, and “positive regulation of cell death” was observed, which reflects cellular senescence and SCD. Notably, we found that genes related to the KEGG pathway “lysosome”, such as *ATP6AP1, CTSH, CTSK, HYAL1, CTSA, PSAP, NPC2, ATP6V0D2, SUMF1*, and genes related to the GO term “hydrogen peroxide metabolic process”, such as *GPX3, HP, MAOB, MT3, PINK1, CHI3L1, CTSH, SLC6A3, IDH3A, ASS1, CRYAB, PDGFRB, ACSL1, GSTM2, IDO1, PCCB, PTGES, PMVK, SLC2A6, CST3, CTSA, DERL3, HCAR2*, were profoundly enriched. These data raised the possibility that alterations in the autophagy-lysosome degradation system and in oxidative stress may have contributed to SCD in NMR cells.

**Figure 3.**
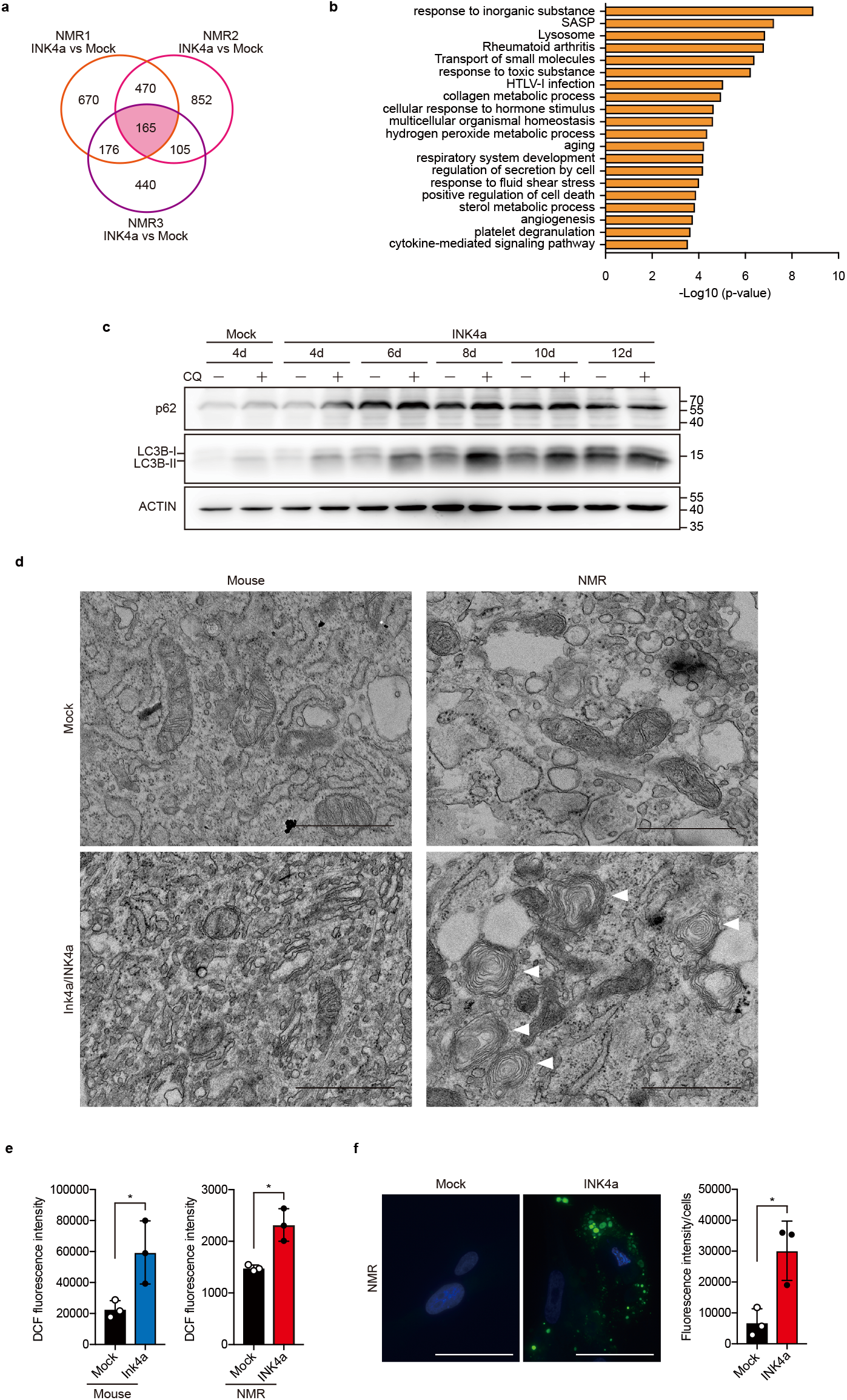
Dysregulation of autophagy and increase in oxidative stress in senescent NMR cells. **a**, Venn diagram showing the DEGs identified from comparisons of mock-transduced and INK4a-transduced NMR-fibroblasts 12 days after transduction. **b**, Top 20 enriched gene ontology (GO) terms and KEGG pathways obtained by using Metascape. **c**, Time-course western blot analysis for LC3B and p62 in chloroquine (CQ)-treated, INK4a-transduced NMR-fibroblasts. ACTIN was used as a loading control. **d**, Electron microscopy images of mock- or INK4a-transduced mouse- or NMR-fibroblasts at 12 days after transduction. Scale bar, 1 μm. **e**, Quantification of ROS using 2’,7’–dichlorofluorescin diacetate (DCFDA) in mock- or INK4a-transduced mouse- or NMR-fibroblasts at 12 days after transduction. **f**, Left, representative images of lipid peroxidation (green) staining in NMR-fibroblasts at 12 days after INK4a-transduction. Scale bar, 50 μm. Right, quantification of signal intensity of lipid peroxidation staining. * *P* < 0.05; *t*-test for **e** and **f**. Data are mean ± SD from *n* = 3 biological replicates.

First, to investigate whether autophagy flux changes in NMR cells during senescence, we performed time-course turnover assay of LC3 and p62, the essential components of autophagosome and selective autophagy respectively, in INK4a-transduced NMR fibroblasts. In this assay, by inhibiting autophagosome-lysosome fusion using chloroquine (CQ), autophagy flux can be detected as LC3-II and p62 protein accumulation levels^46^. On day 8 after INK4a transduction, accumulation of LC3-II with CQ transiently increased, and then the LC3-II level gradually decreased. A remarkable decrease in LC3-II accumulation was observed on day 12, indicating that a transient activation and the subsequent retardation of autophagy flux occurred in NMR fibroblasts undergoing senescence (Fig. 3c). Electron microscopy analysis showed that, in NMR senescent cells on day 12, there were almost no primary lysosomes, an indicator of lysosome biogenesis. On the other hand, an accumulation of multilamellar bodies was observed (white arrowheads in Fig. 3d). Taken together, these results indicate marked dysregulation of the Autophagy-Lysosome system in NMR senescent cells.

Next, we examined whether oxidative stress is increased in NMR fibroblasts during senescence. We measured intracellular ROS level in NMR fibroblasts after INK4a transduction using 2,7-dichlorofluorescin diacetate (DCFH-DA)^47^. A significant increase in the intracellular ROS level (represented by fluorescence intensity) was observed in both senescent NMR and mouse fibroblasts (Fig. 3e). Consistent with this data, senescent NMR cells significantly increased lipid peroxidation level (Fig. 3f). Generally, accumulation of lipid peroxide associates with ferroptosis^48^. However, in senescent NMR-cells, treatment with ferroptosis inhibitors Ferrostatin-1 (Fer-1) and deferoxamine (DFO) did not attenuate SCD, suggesting that ferroptosis is not the major cause of SCD in NMR (Supplementary Fig. 5e). These results demonstrate the dysregulation of autophagy flux and ROS increase in senescent NMR cells.

### Inherent vulnerability to H_2_O_2_ and unique metabolic system in NMR concertedly regulate SCD

Generally, during senescence of human and mouse cells, ROS activation and autophagy impairment are observed^31,32,49^. Nevertheless, unlike senescent NMR cells, senescent mouse and human cells do not die. The key question is why NMR cells induce SCD during senescence. To obtain the mechanistic insight, we focused on the inherent vulnerability of NMR cells. Salmon *et al*. previously revealed that NMR fibroblasts were much more sensitive to treatment with an endoplasmic reticulum (ER) stress inducer and H_2_O_2_ compared to mouse fibroblasts^20^. ER stress and ROS are the well-known stressors which are markedly increased in senescent cells^29^. Thus, we hypothesized that inherent vulnerability to either or both of these stressors might contribute to SCD in senescent NMR cells. Indeed, INK4a-transduction in NMR fibroblasts led to upregulation of apoptosis-inducing ER-stress genes including DNA damage-inducible transcript 3 (DDIT3), also known as C/EBP homologous protein (CHOP), and growth arrest and DNA-damage-inducible 34 (GADD34)^50^ (Supplementary Fig. 5f). However, knockdown of these ER stress genes did not attenuate SCD, indicating that increased ER stress is not the direct cause of SCD (Supplementary Fig. 5g, h).

Next, we found that NMR fibroblasts showed significantly higher vulnerability to treatment with H_2_O_2_ than mouse fibroblasts, consistent with a previous report^20^ (Fig. 4a). This result indicated that NMR fibroblasts have inherent vulnerability to ROS. Generally, upon cellular senescence induction, mammalian cells drastically remodel their metabolism, which is required for their survival and unique phenotype, such as secretion of SASP^24^. Since NMR cells have inherent vulnerability to ROS, we hypothesized that a unique metabolic system in NMR might contribute to SCD. Therefore, we comprehensively measured changes in metabolome of mouse and NMR fibroblasts during senescence. PCA plot showed that, in mouse cells, senescence induction resulted in a large metabolome shift (Fig. 4b). Unexpectedly in NMR cells, the metabolic shift was significantly smaller than that of mouse, while their trend (i.e. contributing loading factor (metabolites)) was similar (Fig. 4b). Volcano plot analysis validated the stable metabolic system against the senescence induction in NMR cells. During senescence induction, several times more metabolites were increased (red zone) or decreased (green zone) in mouse, compared to those in NMR (Fig. 4c). From these results, we wondered that critical metabolic shifts required for SCD induction might be included in this small but concentrated change. Strikingly, we found that two important H_2_O_2_ producing pathways are accelerated in senescent NMR fibroblasts (Fig. 4d, e). The first one is 5-hydroxyindoleacetic acid (5-HIAA) production from serotonin, involving MAO-A activity, which is accompanied by large H_2_O_2_ production^51^. Markedly, serotonin was uniquely accumulated in proliferative NMR fibroblasts, and was converted to 5-HIAA during senescence (Fig. 4d, left). This data indicates the release of large amount of H_2_O_2_ in NMR cells during senescence. We note that the amount of kynurenine, whose production is mediated by Indoleamine 2,3-dioxygenase (IDO), was also significantly increased (Fig. 4d, right). Second pathway is an acceleration of nucleobase degradation involving xanthine oxidase (XO) and uricase activity (Fig. 4e). Since levels of both uric acid and allantoin, downstream metabolites of nucleic acid, were significantly elevated; senescent NMR cells were also exposed to H_2_O_2_ via this pathway (Fig. 4e). It is notable that uricase expression of the liver in NMR was kept uniquely low compared to other mammalian species^52^, thus the acceleration of this enzymatic activity might be another key ROS source for induction of SCD. Finally, to clarify whether the ROS increase contributes to SCD in senescent NMR cells, we treated senescent NMR fibroblasts with N-acetyl L-cysteine (NAC) which acts as an antioxidant in several pathways^53^. The treatment with NAC significantly alleviated SCD in INK4a-transduced senescent NMR fibroblasts (Fig. 4f). Taken together, these results demonstrate that inherent vulnerability to ROS and unique metabolic system in NMR cells concertedly contribute to the accumulation of cellular damage during senescence, and finally inducing SCD in NMR cells (Fig. 4g).

**Figure 4.**
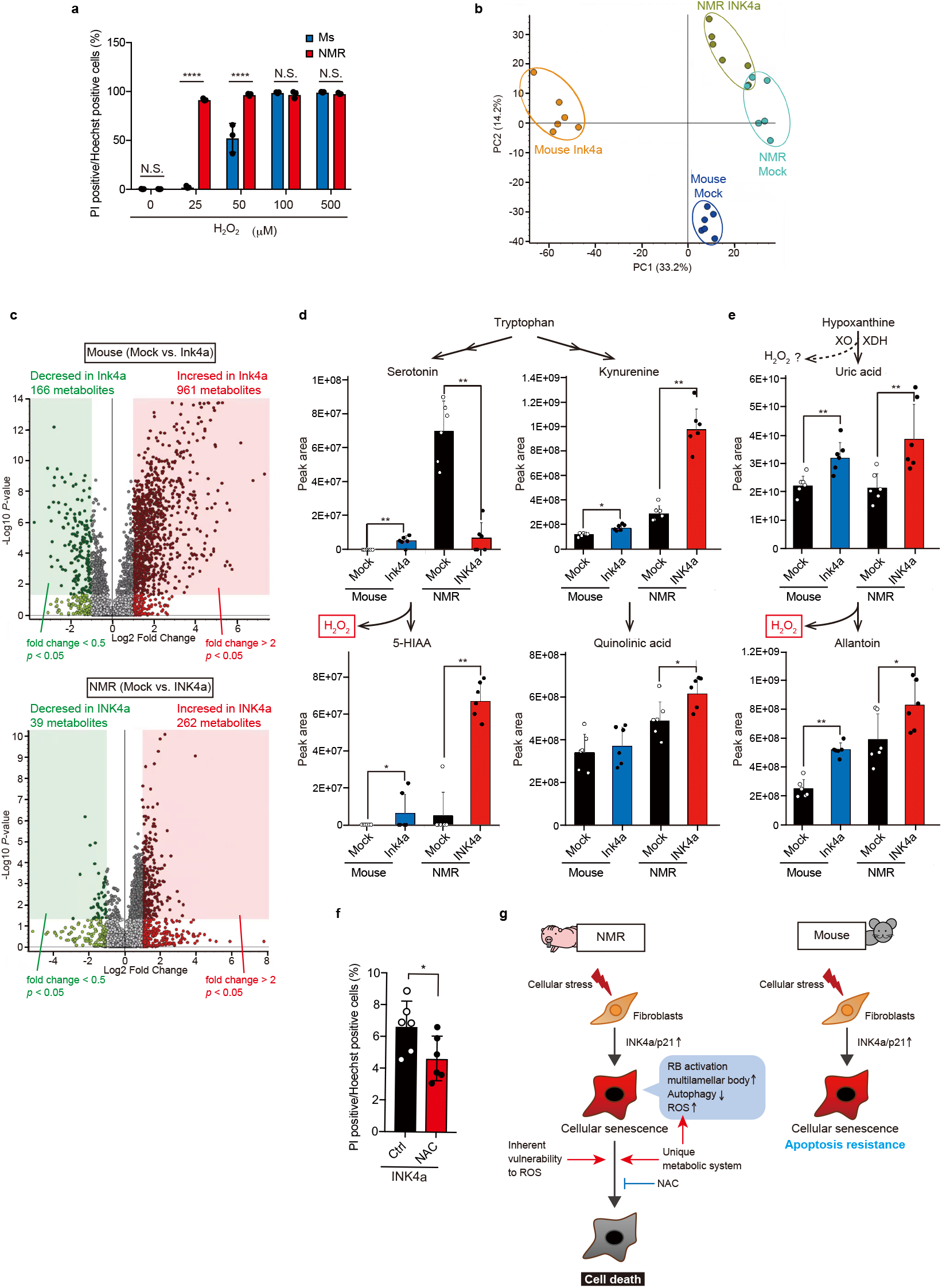
Inherent vulnerability to H_2_O_2_ and unique metabolic system in NMR concertedly regulate SCD. **a**, Quantification of PI-positive cells in NMR-fibroblasts treated with indicated doses of H_2_O_2_ for 6 h. **b**, PCA plot of metabolome differences in mouse- or NMR-fibroblasts at 12 days after INK4a or mock vector transduction. **c**, Volcano plots of metabolome differences in mouse- or NMR-fibroblasts at 12 days after INK4a transduction compared to mock. **d**, Levels of serotonin (upper left), 5-hydroxyindole-3-acetic acid (5-HIAA) (lower left), kynurenine (upper right), and quinolinic acid (lower right) measured by LC-MS/MS in mouse- or NMR-fibroblasts at 12 days after INK4a transduction. Arrows indicate metabolic pathways of each metabolite. Serotonin metabolic pathway can generate H_2_O_2_. Data are mean ± SD from two technical replicates for each cell line (*n* = 3 biological replicates). **e**, Levels of uric acid (upper) and allantoin (lower) measured by LC-MS/MS in mouse- or NMR-fibroblasts at 12 days after INK4a transduction. Arrows indicate metabolic pathways of each metabolite. These pathways can generate H_2_O_2_. XO; xanthine oxidase, XDH; xanthine dehydrogenase. Data are mean ± SD from two technical replicates for each cell line (*n* = 3 biological replicates). **f**, 20 days after INK4a transfection into fibroblasts, the cells were treated with NAC for 24 h and PI-positive cells were quantified (%). Data are mean ± SD from *n* = 6 biological replicates. **g**, Schematic diagram representing SCD in NMR cells and cellular senescence in mouse cells. In NMR cells, inherent vulnerability to ROS and unique metabolic system induces RB-dependent SCD, which simultaneously results in multilamellar bodies accumulation and autophagy dysregulation. * *P* < 0.05; Two-way ANOVA followed by Sidak’s multiple comparisons test for **a**. *t*-test for **d**, **e** and **f.** Data are mean ± SD from *n* = 3 biological replicates except for **d**, **e** and **f**.

## Discussion

In this study, we revealed suppressed accumulation of senescent cells in NMR skin during aging and after UV irradiation *in vivo*. We discovered that NMR fibroblasts exhibit a unique phenomenon, SCD, due to their inherent vulnerability to ROS and unique metabolic system in senescent NMR cells. Recent studies have shown that the accumulation of senescent cells promotes aging of body tissues and a variety of aging-related diseases including cancer, by secreting SASP^54^. Several reports indicated that clearance of senescent cells delays the tissue aging process and aging-related diseases in mice^55^. On the other hand, senescent cells play pivotal roles in homeostatic maintenance, developmental process, and tissue repair^25–27^. Indeed, Grosse *et al*. recently reported that elimination of Ink4a-high senescent cells resulted in liver and perivascular tissue fibrosis^56^. Therefore, there is still controversy over safety of senolytic drugs that kill senescent cells. The discovery of “naturally senolytic” SCD phenotype in the longest-lived rodent NMR supports this concept that the clearing senescent cells during aging would work positively to suppress aging and keep tissues in healthy states. It was reported that NMRs had resistance to various age-related diseases such as Alzheimer’s disease, and cancer^6,57^. SCD may also contribute to resistance against these diseases in NMRs. Therefore, future studies are required to reveal whether other cell types of NMR also induce SCD. Moreover, it is of interest if, after SCD induction, NMR somatic stem cells activate replenishment of new cells in their tissues. Further research is needed to understand the dynamics of senescent cells in other situations such as wound healing in NMRs.

The evolutional adaptation to hypoxic underground environment in NMRs may contribute to longevity via hypometabolism. The evidence of low oxygen consumption rate^58^ and low caloric intake of NMRs^59^ seem to support this theory. Munro *et al*. suggested that the high mitochondrial consumption rate of hydrogen peroxide might be associated with NMR longevity^17^. Perhaps, NMRs may not need to develop a high intracellular anti-oxidant defence capacity because of their hypometabolism and high mitochondrial ROS consuming capacity. Hence, NMRs may be inherently vulnerable to situations with rapid intracellular ROS increase, such as cellular senescence. Interestingly, we identified that NMRs show unique stable metabolic system compared to those in mouse. During senescence, NMR cells showed marked resistance to metabolic remodelling, which is generally important for survival of “damaged” senescent cells. Recently, Zhao *et al*. reported that, during senescence, NMR cells showed smaller change in global gene expression than those in mouse^37^. Considering together, NMR might have an inflexible cellular system that does not respond well to changes in various intracellular situations, such as senescence. Our discovery of SCD explains how vulnerability to ROS and unique metabolic system in this hypometabolic animal species results in the clearance of the damaged senescent cells, which would contribute to delayed aging phenotype. Future studies are required to clarify whether other damaged, non-senescent cells with elevated ROS levels, such as pre-malignant cells, also experience cell death in NMRs. It will also be fascinating to study if cells from other hypoxia-adapted and long-lived animal species, such as blind mole-rats and long-lived bats exhibit similar phenotypes (vulnerability to ROS and inducing SCD) as seen in NMRs.

Zhao *et al*. reported that NMR fibroblasts required a higher dose of irradiation for induction of cellular senescence compared to mouse fibroblasts^37^. Moreover, they found that NMR fibroblasts exhibited a lower proportion of irradiation-induced acute cell death at day 3 after irradiation than mouse fibroblasts, which occurs prior to senescence. This phenotype would be due to the efficient double-strand break repair system in NMRs as previously reported by Tian *et al*^60^. In this study, we utilized INK4a transduction method to induce only cellular senescence but not acute cell death, and discovered the unique SCD phenotype in NMRs. Considering these results together, NMR cells likely have a double safeguard system to inhibit the accumulation of senescent cells: efficient DNA double-strand break repair and SCD.

Recently, Takasugi *et al*. reported that extracellular very-high-molecular-mass hyaluronan (vHMM-HA) in NMR had cytoprotective properties compared to the shorter HMM-HA^61^. In NMR fibroblasts, degradation of vHMM-HA by HAase decreased the viability upon oxidative stressor, tBHP-treatment. This report and our results suggest that NMR may protect their body from stresses by several unique mechanisms: NMR cells are protected from extracellular stress by vHMM-HA and when accumulation of intracellular oxidative stress exceeds a certain level upon senescence, NMR cells go into SCD via activation of Rb pathway. We speculate that previously reported cell death tolerance of NMR vessels to H_2_O_2_ might be due to the existence of extracellular vHMM-HA in tissues or due to difference in cell types^18^.

Our study proposes that SCD would contribute to maintaining the quality of cells in tissues by eliminating “damaged” senescent cells in NMRs. In other long-lived and cancer-resistant mammalian species, some species-specific systems which induce cell death in “damaged” cells were reported. In elephant, there are 40 copies of the tumour suppressor p53 gene, and their lymphocytes are more sensitive to irradiation than human lymphocytes, allowing only cells with lower DNA damage to survive^62^. Fibroblasts from blind mole rats are known to induce species-specific, concerted necrotic cell death when cells are stressed by *in vitro* long culture^63^. It is of great interest that several different systems for maintaining the quality of cells in tissues may have evolved in the long-lived and cancer-resistant animal species. Further studies are required to reveal how and how much long-lived animals can keep the quality of cells and remove “damaged” cells in tissues.

In this study, we discovered that NMRs had a unique SCD phenotype, most likely associated with adaptation to underground hypoxic conditions and their delayed aging phenotype. Thus, advance studies in NMRs are useful to provide molecular and mechanistic clues to maintain human bodies healthier and long-living. Subsequent further research on detailed molecular mechanisms underlying SCD in NMRs would contribute to development of safer senolytic drugs to delay aging and understanding of age-related diseases in humans.

## Methods

### Animals

The Ethics Committees of Kumamoto University (approval no. A30-043 and A2020-042), Hokkaido University (approval no. 14-0065) and Keio University (approval no. 12024) approved all procedures, which were in accordance with the Guide for the Care and Use of Laboratory Animals (United States National Institutes of Health, Bethesda, MD). NMRs were maintained in Kumamoto University. C57BL/6 mice and CD1 mice were purchased from CLEA Japan, Inc. HR1 mice were purchased from Japan SLC, Inc. Cells and tissues were obtained from at least three animals.

### In vivo analysis

To obtain skin samples without sacrifice, mice [CD1; young (4-week old) and middle-aged (one-year old] and NMRs [young (one-year old) and middle-aged (13-15-year old, the current available oldest animals in our laboratory)] were anesthetized with isoflurane (FUJIFILM Wako), and four 3 mm-diameter biopsy punches (three animals per experimental group) were made on the dorsal surfaces. Skin biopsy samples were subjected to SA-β-gal staining and RNA isolation. To induce cellular senescence in the skin, HR1 mice (6-week old) and NMRs (one-year old) were exposed to 1000 J/m^2^ of UV-B using a UV lamp (UVP UVM-28, analytikjena) for indicated times (seven exposures in two weeks, as shown in Supplementary Fig. 1b, three animals per experimental group). Irradiated skins were excised and subjected to histological analysis and RNA extraction. Due to the limitation of usable number of NMRs (deriving from low breeding rate of NMRs), three NMRs were used in *in vivo* experiments (which is the number used in many NMR studies).

### Cell culture and drug treatment

Primary NMR or mouse fibroblasts were isolated from back skin or lung of 1–2-years-old adult NMRs or 6-week-old adult C57BL/6 mice. Primary fibroblasts prepared from tissues of at least three different animals were used as biological replicates. Tissues were washed with ice-cold phosphate-buffered saline (PBS; Nacalai tesque) containing 1% penicillin/streptomycin (FUJIFILM Wako) and amphotericin B (FUJIFILM Wako). Tissues were then minced, and then suspended in the culture medium (contents described below), plated on gelatine-coated 10-cm cell culture dishes (IWAKI), and cultured at 32°C in a humidified atmosphere containing 5% CO_2_, 5% O_2_. Cells were cultured in the medium composed of Dulbecco’s Modified Eagle’s Medium (DMEM, Sigma) supplemented with 15% foetal bovine serum (FBS) (for NMR-fibroblasts) or 10% FBS (for mouse-fibroblasts) (BioWest or Gibco, at least three lots), 1% penicillin/streptomycin, 2 mM L-glutamine (Nacalai tesque or FUJIFILM Wako) and 0.1 mM non-essential amino acids (NEAA, Nacalai tesque or FUJIFILM Wako). Fibroblasts were used within 5 passages. Medium was replaced every 2 days. A 10 mM stock solution of chloroquine diphosphate (CQ; Sigma) was prepared with water (Sigma) and filtered with 0.22 μm syringe filter (Sartorius), and used at 10 μM. A 1 M stock solution of NAC (FUJIFILM Wako) was prepared in 20 mM HEPES buffer (Nacalai tesque), titrated with NaOH to pH 7.4, and filter sterilized and used at 0.5 mM. Z-VAD-FMK was dissolved in dimethyl sulfoxide (DMSO; Sigma) at 20 mM as a stock solution and used at 20 μM. HAase was purchased from Sigma and used at 3 U/ml. Ferrostain-1 (Fer-1) and deforoxamine (DFO) were kindly provided by Dr. Toshiro Moroishi (Kumamoto University). CQ, NAC, Z-VAD-FMK and HAase were added during the 24 h period before cell death analysis. Fer-1 and DFO were added for 2 h. All the experiments were performed in triplicates.

### Lentivirus preparation and infection

We used lentiviral vectors pCSII-EF-NMR-INK4a-TK-hyg, pCSII-EF-NMR-INK4a, pCSII-EF-mouse-Ink4a-TK-hyg, pCSII-EF-mouse-Ink4a, pCSII-EF-HRasV12-TK-hyg, pCSII-EF-NMR-p15/p21/p27/pALT-TK-hyg, pCSII-EF-human-BCL-XL-TK-hyg or pCSII-EF-NMR-BCL-XL-TK-hyg for ectopic expression and H1 promoter-driven vectors previously described for shRNA expression^64^. The backbone vectors for ectopic expression (pCSII-EF-RfA) and shRNA expression were purchased from RIKEN BioResource Research Center. The backbone vector for ectopic expression (pCSII-EF-RfA-TK-Hyg) was kindly provided by Dr. Hayato Naka-Kaneda (Shiga University of Medical Science). The knockdown vectors expressing shRNA against NMR-INK4a^65^, NMR-DDIT3 or NMR-GADD34 were generated to target the sequences shown in Supplementary Table 2. Each plasmid and packaging vectors (pCMV-VSV-G-RSV-Rev and pCAG-HIVgp) were used to transfect HEK293T cells with Polyethylenimine MAX transfection reagent (Polysciences), according to the manufacturer’s instructions. Nine hours after transfection, the medium was replaced, the conditioned medium containing viral particles was collected two times every 24 hr.

For lentiviral infection, cells were seeded at 3 × 10^5^ cells/10-cm dish one day before infection. The conditioned medium containing lentivirus was filtered with 0.45 μm syringe filter (Sartorius) and diluted two-fold in growth medium, and was used for viral transduction. 24 hr after the first infection, the medium was replaced with second conditioned medium containing lentivirus. After viral infection, the medium was replaced with growth medium, which was changed every 2 days.

### SA-β-Gal activity analysis

For measuring cellular senescence, the SA-β-Gal staining was performed using Senescence Detection Kit (BioVision). Cells or fresh-frozen skin sections were stained according to manufacturer’s instructions for 48 h at 37 °C or 32 °C. The cells or skin sections were washed with PBS and stained with Hoechst 33258 (1:1000 dilution in PBS) for 10 minutes at room temperature in the dark. The cells or skin sections were washed with PBS and analysed using microscopy (Keyence). Entire cell populations in three random microscope fields (at least 150 cells) were analysed for perinuclear blue staining indicative of SA-β-gal activity and Hoechst positive nuclei. To analyse skin sections, at least three random microscope fields from three animals per experiment were obtained. Hair follicle regions were excluded from quantitative analysis because these regions are constitutively positive for SA-β-Gal activity regardless of cellular senescence. SA-β-Gal positive regions were quantified by ImageJ. To stain floating cells, the culture supernatant was spun down, and the cell pellet was resuspended in Smear Gell kit (GenoStaff) and spread on the slide surface. The slides were stained and analysed in the same way as adherent cells.

### Quantitative RT-PCR

Mouse and NMR fibroblasts were harvested and washed with ice-cold PBS. Cells were spun down, and the pellet was used for isolation of total RNA using the RNeasy Plus Mini Kit (Qiagen). To remove genomic DNA, gDNA Eliminator spin columns were used. RNA was eluted from the columns using 30 μl of RNase-free water and quantified using a NanoDrop spectrophotometer (Thermo Scientific). cDNA was synthesized with the ReverTra Ace qPCR RT Master Mix (TOYOBO) using 400 ng of total RNA input. Real-time quantitative PCR reactions were set up in triplicate using SYBR Premix Ex Taq™ II (Tli RNaseH Plus) (TaKaRa), Fast SYBR Green Master Mix (Invitrogen) or THUNDERBIRD SYBR qPCR Mix (TOYOBO), and run on a ViiA 7, StepOne plus Real-Time PCR System (Applied Biosystems) or CFX384 Touch Real-Time PCR Detection System (Bio-Rad). Primer sequences are listed in Supplementary Table 3.

### Immunohistochemistry

Fresh-frozen skin sections (10 μm) were fixed with 4% PFA for 10 min at room temperature, washed with PBS and then blocked with 5% normal goat serum in PBS for 60 min at room temperature. The sections were incubated with primary antibodies against cleaved caspase 3 (CST; 9664; 1:400) in Can Get Signal Solution B (TOYOBO) for 12 h at 4°C. After washes with PBS, the sections were incubated with secondary antibody Alexa Fluor 555 anti-rabbit IgG (CST; A21429; 1:1000), and nuclei were counterstained with 1 μg /ml Hoechst 33258 (Sigma–Aldrich) for 60 min at room temperature. After washes with PBS, images were captured. TUNEL staining (for quantifying cell death marker) was performed using TUNEL Assay Kit BrdU-Red (abcam; ab66110) according to the manufacturer’s instructions. The images were captured by BZ-X 710 fluorescence microscope (Keyence) and analysed using a BZ-X image analyzer (Keyence). To analyse skin sections, images from at least three random microscope fields from three animals per experiment were obtained.

### 5-bromo-2-deoxyuridine (BrdU) incorporation assay

To analyse cell proliferation, BrdU labelling was performed for 2 days for mouse fibroblasts, and 4 days for NMR fibroblasts as previously described^7^. Then, cells were fixed with 4% paraformaldehyde (PFA; FUJIFILM Wako) in PBS and subjected to immunostaining. BrdU was detected using primary sheep antibody against BrdU (Fitzgerald; 20-BS17; 1:200) and Alexa Fluor 555 anti-sheep IgG (A11015; Life Technologies; 1:500) secondary antibody. Cell nuclei were stained with 1 μg /ml Hoechst 33258 (Sigma-Aldrich). Cells were observed under a BZ-X 710 fluorescence microscope (Keyence) and counted using a BZ-X image analyzer (Keyence). Entire cell populations in four random microscope fields (at least 150 cells) per 3 cell lines were analysed for BrdU positive and Hoechst positive nuclei.

### Flow cytometry analysis for cell death detection

Cell death was examined using FITC Annexin V Apoptosis Detection kit (BD Biosciences or BioLegend). Adherent cells were harvested, stained according to manufacturer’s protocols. Flow cytometry was performed with a FACSCalibur or FACSVerse (BD Biosciences) flow cytometer. The data were analysed using FlowJo 10 software (BD Biosciences). The experiments were performed in triplicates.

### Hoechst-propidium iodide (PI) staining assay

Cells were seeded into 24-well plate or 60-mm dish and stained with Hoechst 33342 (DOJINDO; 1 mg/ml; 1:1000 in growth medium) for 10 min at 32°C. Then the cells were stained with PI (10 mg/ml; FUJIFILM Wako; 1:1000 in growth medium) for 5 min at 32°C. The images were captured by BZ-X 710 fluorescence microscope (Keyence), and positive cells were counted using a BZ-X image analyzer (Keyence). Entire cell populations in 8-12 random microscope fields (at least 350 cells) per 3 cell lines were analysed for PI positive and Hoechst positive nuclei.

### Western blotting

The cells were washed with PBS, lysed in cell-lysis buffer (62.5 mM Tris-HCl, pH 6.8, 2% SDS and 5% sucrose) and boiled for 5 min. Protein concentrations were measured using BCA Protein Assay Kit (TaKaRa). The samples were subjected to SDS-PAGE, and transferred to a PVDF membrane using a Trans-Blot Turbo Transfer System (Bio-Rad). Membranes were probed with antibodies against NMR-INK4a (non-commercial^65^; 1:1000), AKT (CST; 9272; 1:1000), pAKT (CST; 4060; 1:1000), RB (CST; 9309; 1:1000 for NMR, CST; 9313; 1:1000 for mouse), pRB (CST; 8516; 1:1000), LC3B (CST; 2775; 1:1000) and β-Actin (CST; 4970; 1:2000). The membranes were incubated with HRP-conjugated anti-rabbit (CST; 7074; 1:1000) or anti-mouse (CST; 7076; 1:1000) IgG secondary antibodies and visualized using ECL Western Blotting Detection System or ECL Prime Western Blotting Detection Reagent (Amersham). LAS-4000mini imaging system (FUJIFILM) was used for signal detection and Multi Gauge V3.0 software (FUJIFILM) was used for data analysis. The experiments were performed in triplicates. Uncropped images of all the blots are shown in Source data file.

### Measurement of Intracellular ROS

For detection of cellular ROS, DCFDA/H2DCFDA Cellular ROS Assay Kit (abcam; ab113851) was used according to manufacturer’s protocol. Briefly, fibroblasts were plated on a black 96-well plate with clear bottom for overnight. Cells were treated with 25 μM of DCFDA solution for 45 min at 32°C in the dark. After washing once with PBS, the plate was subjected to a fluorescence plate reader (GloMax; Promega) at Ex/Em = 485/535 nm. The experiments were performed in triplicate.

### Detection of lipid peroxidation

For detection of lipid peroxidation, Click-iT lipid peroxidation imaging kit (Invitrogen) was used according to manufacturer’s protocol. Briefly, cells were plated on coverslips in a 24-well plate and treated with Click-it linoleamide alkyne (LAA) for 24 h at 32 °C. After washing with PBS, cells were fixed in 4% PFA for 15 min at room temperature, washed with PBS, permeabilized with 0.05% TritonX-100 in PBS for 10 min, and blocked with 1% BSA in PBS for 30 min. Cells were washed, and the click reaction was performed with 5 μM Alexa Fluor 488 azide for 30 min. After washing, cells were stained with Hoechst 33258. The images were captured by BZ-X 710 fluorescence microscope (Keyence) and analysed using a BZ-X image analyzer (Keyence). Entire cell populations in 12 random microscope fields (at least 150 cells) per 3 cell lines were analysed for green fluorescence and Hoechst positive nuclei.

### H_2_O_2_ sensitivity test

Cells in a 24-well plate were incubated with indicated doses of H_2_O_2_ (Nacalai tesque) for 6 h in DMEM. Cells were then washed and incubated with growth medium for 18 h. To evaluate cell death, Hoechst-PI staining assay was performed. Entire cell populations in 6 random microscope fields (at least 150 cells) per 3 cell lines were analysed for PI positive and Hoechst positive nuclei.

### RNA-sequencing

NMR fibroblasts were homogenized in TRIzol reagent (Invitrogen). RNA extraction and library preparation were performed at Novogene Bioinformatics Institute. NMR samples were sequenced using the Novaseq 6000 (150 bp paired-end). NMR reference genome assemblies (GRCm38 and HetGla_female_1.0) and corresponding annotation files were obtained from Ensembl release 92^66^. Raw reads were trimmed using Trim Galore (ver. 0.5.0, https://www.bioinformatics.babraham.ac.uk/projects/trim_galore/), and the transcript abundances (transcripts per million [TPM]) were calculated using RSEM (ver. 1.2.25)^67^ with Bowtie 2^68^ (ver.2.2.6).

### Metabolome analysis

Metabolite extraction from cultured cells was performed as described previously^69^. Briefly, frozen cells were lysed and scraped with ice-cold methanol (500 μl) together with internal standard (IS) compounds (see below), followed by the addition of an equal volume of ultrapure water and 0.4 times the volume of chloroform (LC/MS grade, FUJIFILM Wako). The suspension was then centrifuged at 15,000 g for 15 min at 4°C. After centrifugation, the aqueous phase was ultrafiltered using an ultrafiltration tube (Ultrafree MC-PLHCC, Human Metabolome Technologies). The filtrate was concentrated with a vacuum concentrator (SpeedVac, Thermo). The concentrated filtrate was dissolved in 50 μl of ultrapure water and used for LC-MS/MS and IC-MS analyses. As, internal standard (IS) compounds, we used 2-morpholinoethanesulfonic acid (MES) and L-methionine sulfone as ISs for anionic and cationic metabolites, respectively. These compounds are not present in the tissues; thus, they serve as ideal standards. Loss of endogenous metabolites during sample preparation was corrected by calculating the recovery rate (%) for each sample measurement.

For metabolome analysis, anionic metabolites were measured using an orbitrap-type MS (Q-Exactive focus, Thermo Fisher Scientific, San Jose, CA), connected to a high performance ion-chromatography system (ICS-5000+, Thermo Fisher Scientific) that enables us to perform highly selective and sensitive metabolite quantification owing to the IC-separation and Fourier Transfer MS principle. The IC was equipped with an anion electrolytic suppressor (Thermo Scientific Dionex AERS 500) to convert the potassium hydroxide gradient into pure water before the sample enters the mass spectrometer. The separation was performed using a Thermo Scientific Dionex IonPac AS11-HC, 4-μm particle size column. IC flow rate was 0.25 ml/min supplemented post-column with 0.18 ml/min makeup flow of MeOH. The potassium hydroxide gradient conditions for IC separation are as follows: from 1 mM to 100 mM (0–40 min), 100 mM (40–50 min), and 1 mM (50.1–60 min), at a column temperature of 30°C. The Q Exactive focus mass spectrometer was operated under an ESI negative mode for all detections. Full mass scan (m/z 70-900) was used at a resolution of 70,000. The automatic gain control (AGC) target was set at 3 × 106 ions, and maximum ion injection time (IT) was 100 ms. Source ionization parameters were optimized with the spray voltage at 3 kV and other parameters were as follows: transfer temperature at 320°C, S-Lens level at 50, heater temperature at 300°C, Sheath gas at 36, and Aux gas at 10.

The amounts of cationic metabolites (amino acids) was quantified using liquid chromatography-tandem mass spectrometry (LC-MS/MS). Briefly, a triple-quadrupole mass spectrometer equipped with an ESI ion source (LCMS-8060, Shimadzu Corporation) was used in the positive and negative-ESI and multiple reaction monitoring (MRM) modes. The samples were resolved on the Discovery HS F5-3 column (2.1 mmI.D. x 150 mmL, 3 μm particle, Sigma-Aldrich), using a step gradient with mobile phase A (0.1% formate) and mobile phase B (0.1% acetonitrile) at ratios of 100:0 (0–5 min), 75:25 (5–11 min), 65:35 (11–15 min), 5:95 (15–20 min), and 100:0 (20–25 min), at a flow rate of 0.25 ml/min and a column temperature of 40°C. MRM conditions for each amino acids were previously described^70^.

### Transmission electron microscopy

Cells were immediately fixed in 2% glutaraldehyde (TAAB)/2% PFA (FUJIFILM Wako)/ 30 mM HEPES buffer for 30 min at room temperature. After post-fixation with 1% OsO4 (Merck) and en bloc staining with 1.5% uranyl acetate, cells were embedded in Araldite (TAAB) and examined in a transmission electron microscope (Hitachi H-7700).

### Mitomycin C treatment

Mouse and NMR fibroblasts were exposed twice to mitomycin C (MMC) at 200 nM for 24 hours. MMC-containing medium was added to subconfluent fibroblasts. After 24 h, the medium was replaced by a freshly prepared MMC-containing medium for additional 24 h. Then, the cells were washed and cultured in fresh medium for 10 days.

### Statistical analysis

Prism 7 software (GraphPad) was used for statistical analysis. Data were presented as the mean ± standard deviation (SD). Data were analysed using two-way ANOVA followed by Sidak’s multiple comparisons test or one-way ANOVA with Tukey’s multiple comparison test or Dunnett’s multiple comparison test. Unpaired t-tests were used to compare the two groups.

## Acknowledgements

We thank Drs. K. Tomizawa, T. Chujo, J. Kohyama, K. Seino, H. Wada, M. Nakanishi, Y Johmura, E. Hara for their administrative support and scientific discussion, M. Kobe, Y. Tanabe, Y. Fujimura, N. Arai, C. Fukaya for their help with animal maintenance, Y. Tanoue, Y. Takahashi for electron microscopy analysis and all members of the K.M. laboratories for technical assistance and scientific discussion, T. Moroishi for ferroptosis inhibitors, H. Miyoshi for lentiviral vectors, and H. Naka-Kaneda for pCSII-EF-RfA-TK-hyg vector. We thank to Jane Doe of the Liaison Laboratory Research Promotion Center for technical support. We thank Enago (www.enago.com) for the manuscript review and editing support. This work was supported in part by AMED under Grant Number JP20gm5010001, 19bm0704040, and PRESTO of the Japan Science and Technology Agency to K.M., Grants-in-Aid for Scientific Research from the Japanese Society for the Promotion of Science from the Ministry of Education, Culture, Sports, Science and Technology (MEXT) to K.M. and Y.K., Tenure-Track Grant of Kumamoto University to Y.K. K.M. was supported by the Takeda Science Foundation, Mitsubishi Foundation, Japan Foundation For Aging And Health, KOSÉ Cosmetology Research Foundation, The Nakatomi Foundation, and Naito Foundation. This work was also supported, in part, by ROIS-DS-JOINT (012RP2018) to K.M.

## Author contributions

Y.K. conducted most of the experiments; K.O., M.T., Y.O., S.F., S.H. and S.M. conducted certain experiments; H.B., M.T. and Y.K conducted RNA-seq; Y.S. conducted liquid chromatography-mass spectrometry analysis; T.F., M.S., M.N. and H.O. provided technical support in this study; Y.K. and K.M. designed the study; Y.K., H.O., H.B., Y.S., T.F., M.S. and K.M. wrote the manuscript; H.O. and K.M. supervised the project.

## Data availability

RNA-seq data is deposited in the DDBJ under accession number DRA009592.

## Ethics declarations

The authors declare that there are no competing financial interests.

## Supplementary Tables

Supplementary Tables are provided as a Source data file.

---

**Supplementary Figure 1.**
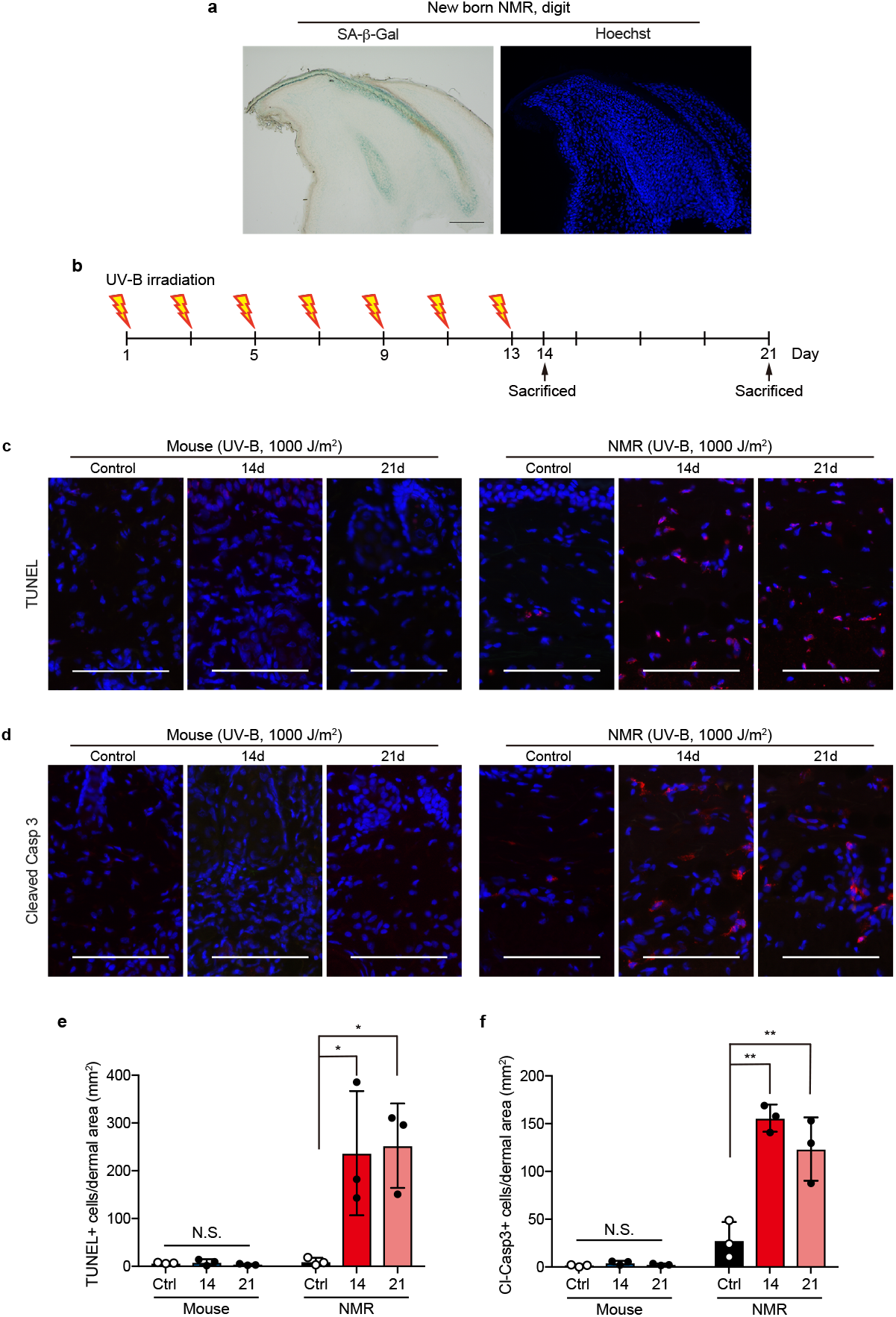
Dead cells but not senescent cells are increased in NMR skin by UV irradiation. **a**, SA-β-Gal staining of neonate NMR digits (Postnatal day 0) for positive control of SA-β-Gal activity in NMR tissue. Scale bar, 100 μm. **b**, Scheme for UV-B irradiation to the skin. **c**, TUNEL staining of skins (dermis) after UV-B irradiation. Scale bar, 100 μm. **d**, Immunohistochemistry for cleaved-Caspase3 in skins (dermis) after UV-B irradiation. Scale bar, 100 μm. **e**, Quantification of TUNEL-positive cells (%) in the dermal area after UV-B irradiation. **f**, Quantification of cleaved Caspase3-positive cells (%) in the dermal area after UV-B irradiation. * *P* < 0.05; ** *P* < 0.01; N.S., not significant, one-way ANOVA followed by Dunnett’s multiple comparison test for e and f. Data are mean ± SD from *n* = 3 animals.

**Supplementary Figure 2.**
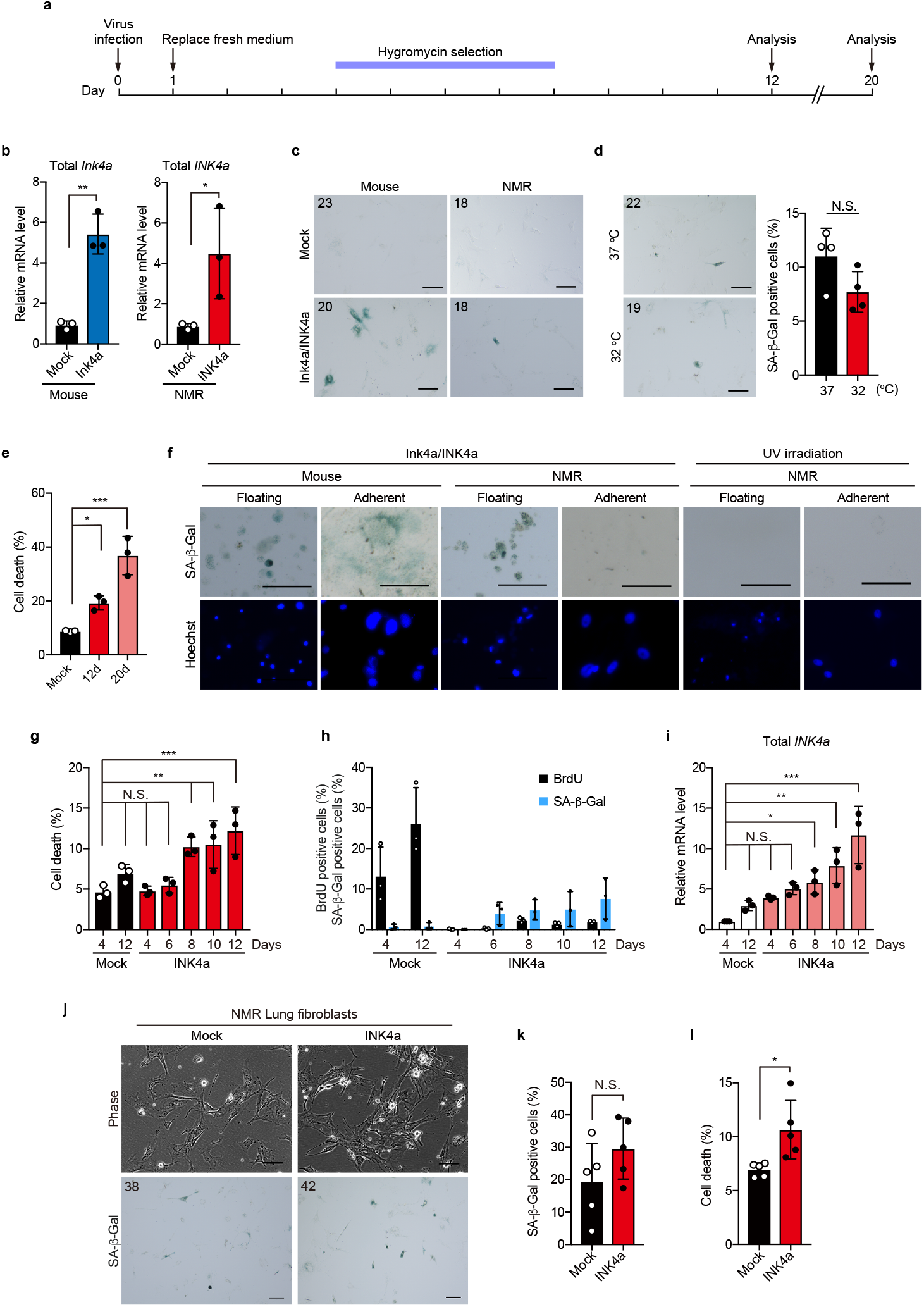
Induction of cellular senescence by INK4a transduction. **a**, Scheme for cellular senescence induction by transduction of INK4a. **b**, qRT-PCR analysis of total INK4a expression in mouse- or NMR-skin fibroblasts. *n* = 3 biological replicates. **c**, SA-β-Gal activity of mouse- or NMR-skin fibroblasts at 12 days after INK4a transduction. The number in the upper left corner indicates Hoechst-positive nuclei. Scale bar, 100 μm. **d**, Comparison of SA-β-Gal activity of NMR-fibroblasts at 37 °C and 32 °C. *n* = 4 biological replicates. The number in the upper left corner indicates Hoechst-positive nuclei. Scale bar, 100 μm. **e**, Quantification of Annexin V-positive cells (%) in INK4-transduced NMR-fibroblasts at 12 days or 20 days after transduction. *n* = 3 biological replicates. **f**, SA-β-Gal activity of floating dead cells and adherent living cells in NMR-fibroblast culture at 12 days after INK4a transduction. Scale bar, 100 μm. NMR-fibroblasts treated with high-dose UV-C (2000 J/m^2^) were used as a control induced acute cell death. **g-i**, Time-course analysis of NMR-fibroblasts after INK4a transduction: quantification of Annexin V-positive cells (%) (**g**); quantification of BrdU- and SA-β-Gal-positive cells (%) (**h**); qRT-PCR for INK4a expression (**i**). *n* = 3 biological replicates. **j**, Cell morphology and SA-β-Gal activity of NMR-lung fibroblasts at 12 days after INK4a transduction. Scale bar, 100 μm. The number in the upper left corner indicates Hoechst-positive nuclei. **k**, Quantification of SA-β-Gal-positive cells in NMR-lung fibroblasts at 12 days after INK4a transduction (%). **l**, Quantification of Annexin V-positive cells in NMR-lung fibroblasts at 12 days after INK4a transduction (%). *n* = 5 biological replicates. * *P* < 0.05; ** *P* < 0.01; *** *P* < 0.001; *t*-test for **b**, **d**, **k** and **l**; one-way ANOVA followed by Dunnett’s multiple comparison test for **e**, **g** and **i**. Data are mean ± SD.

**Supplementary Figure 3.**
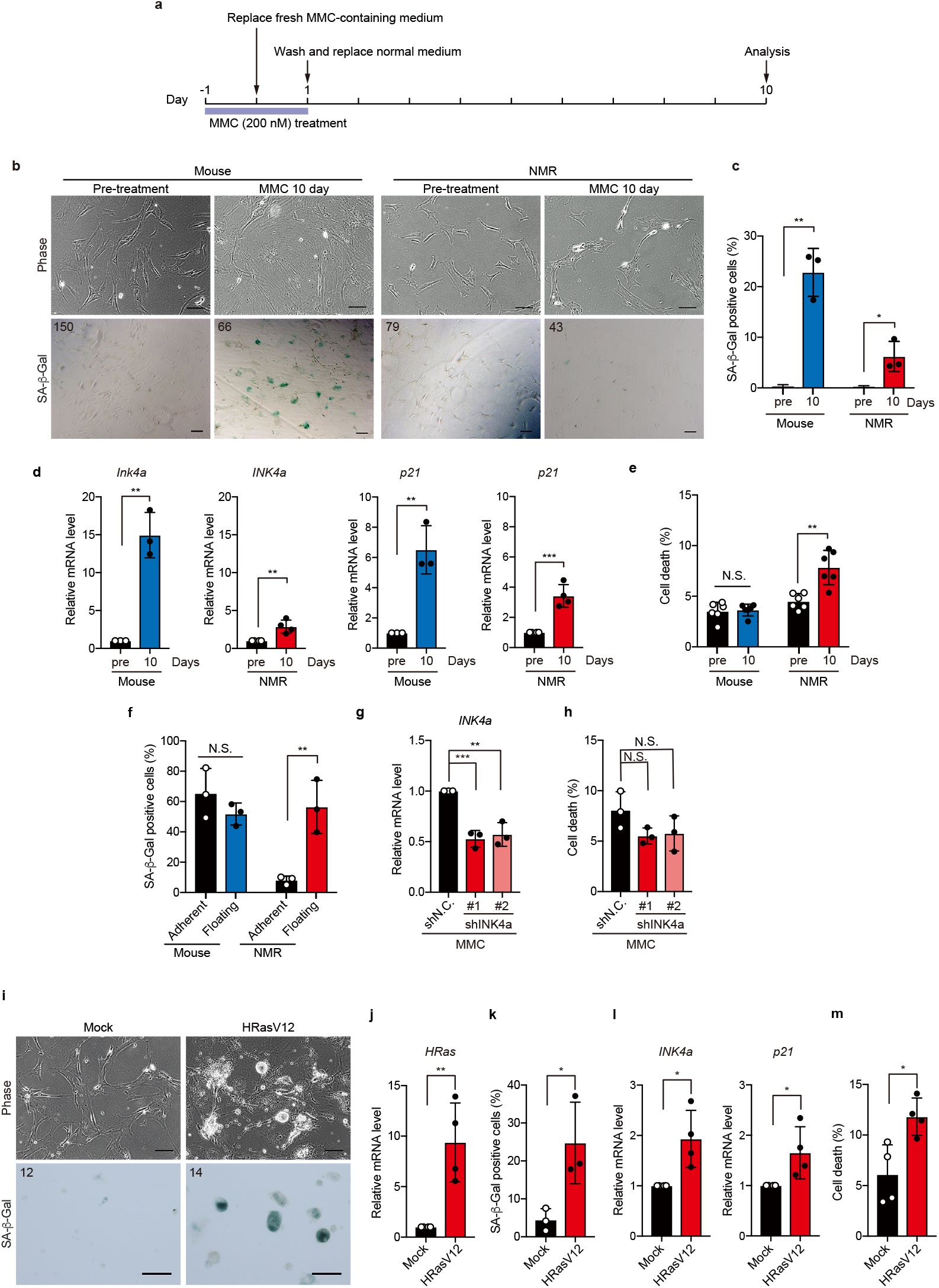
Induction of DNA damage-induced senescence or oncogene-induced senescence evokes SCD in NMR-fibroblasts. **a**, Scheme for cellular senescence induction by low concentration of MMC. **b**, Cell morphology and SA-β-Gal staining. Scale bar, 100 μm. The number in the upper left corner indicates Hoechst-positive nuclei. **c**, Quantification of SA-β-Gal-positive cells (%) in mouse- or NMR-fibroblasts 10 days after MMC treatment. **d**, qRT-PCR analysis for the expression of *INK4a* and *p21* in mouse- or NMR-fibroblasts 10 days after MMC treatment. **e**, Quantification of Annexin V-positive cells (%). *n* = 6 biological replicates. **f**, Quantification of SA-β-Gal-positive cells in floating dead cell population and adherent living cell population in NMR-fibroblast culture at 10 days after MMC treatment (%). **g**, qRT-PCR analysis for the expression of *INK4a* in shINK4a-transduced NMR-fibroblasts at 10 days after the MMC treatment, and quantification of Annexin V-positive cells (%) (**h**). **i**, Cell morphology and SA-β-Gal activity of NMR-fibroblasts 29 days after HRASV12 transduction. Scale bar, 100 μm. The number in the upper left corner indicates Hoechst-positive nuclei. **j**, qRT-PCR analysis for the expression of *HRas*. **k**, Quantification of SA-β-Gal-positive cells (%). **l**, qRT-PCR analysis for the expression of *INK4a* and *p21*. **m**, Quantification of Annexin V-positive cells (%). * *P* < 0.05; ** *P* < 0.01; *** *P* < 0.001; *t*-test for **c, d, e, f, j, k, l** and **m**; one-way ANOVA followed by Dunnett’s multiple comparison test for **g** and **h**. Data are mean ± SD from *n* = 3 biological replicates except for e (*n* = 6), **j, l** and **m** (*n* = 4).

**Supplementary Figure 4.**
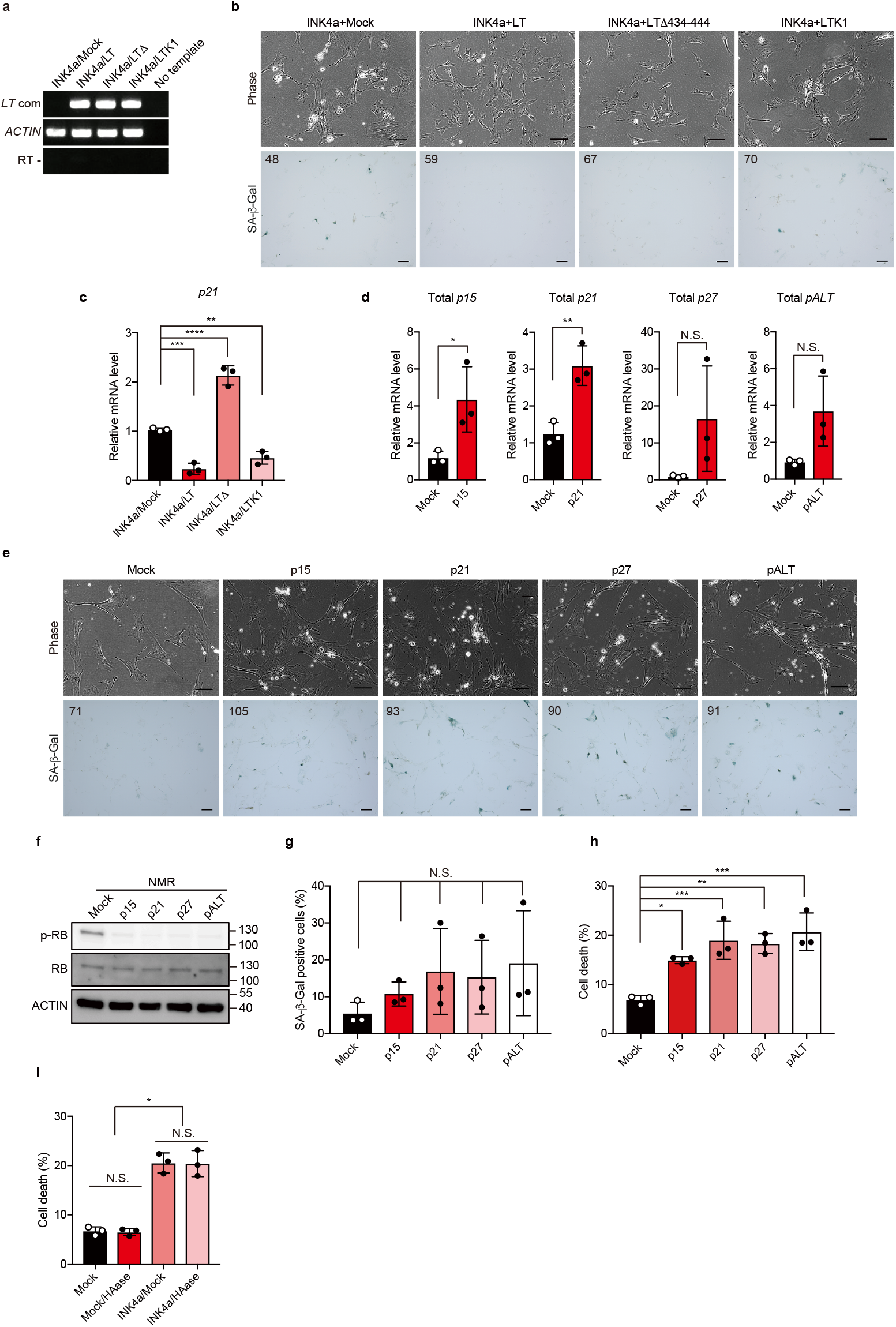
RB but not p53 pathway is essential for the induction of SCD. **a**, RT-PCR analysis of *SV40 Large T* expression in NMR-fibroblasts transduced with different forms of SV40 Large T antigen (LT, LTΔ434–444, and LTK1) and INK4a. **b**, Cell morphology and SA-β-Gal staining of NMR-fibroblasts transduced with different forms of SV40 Large T antigen (LT, LTΔ434–444, and LTK1) and INK4a. Scale bar, 100 μm. The number in the upper left corner indicates Hoechst-positive nuclei. **c**, qRT-PCR analysis for the expression of *p21* in NMR-fibroblasts transduced with different forms of SV40 Large T antigen (LT, LTΔ434–444, and LTK1) and INK4a. **d**, qRT-PCR analysis for the expression of each CKI (*p15,p21,p27* and *pALT*) in NMR-fibroblasts transduced with each CKI. **e**, Cell morphology and SA-β-Gal staining of NMR-fibroblasts transduced with each CKI. Scale bar, 100 μm. The number in the upper left corner indicates Hoechst-positive nuclei. **f**, Western blotting of RB and phospho-RB (p-RB) protein in NMR-fibroblasts transduced with each CKI (p15, p21, p27 and pALT). ACTIN was used as a loading control. **g**, Quantification of SA-β-Gal-positive cells (%). **h**, Quantification of Annexin V-positive cells (%) in NMR-fibroblasts transduced with each CKI. **i**, Quantification of Annexin V-positive cells (%) in INK4a-overexpressed NMR-fibroblasts with HAase treatment. * *P* < 0.05; ** *P* < 0.01; *** *P* < 0.001; **** *P* < 0.0001; *t*-test for **d**; one-way ANOVA followed by Dunnett’s multiple comparison test for **c, g** and **h**, and one-way ANOVA followed by Tukey’s multiple comparison test for **i**. Data are mean ± SD from *n* = 3 biological replicates.

**Supplementary Figure 5.**
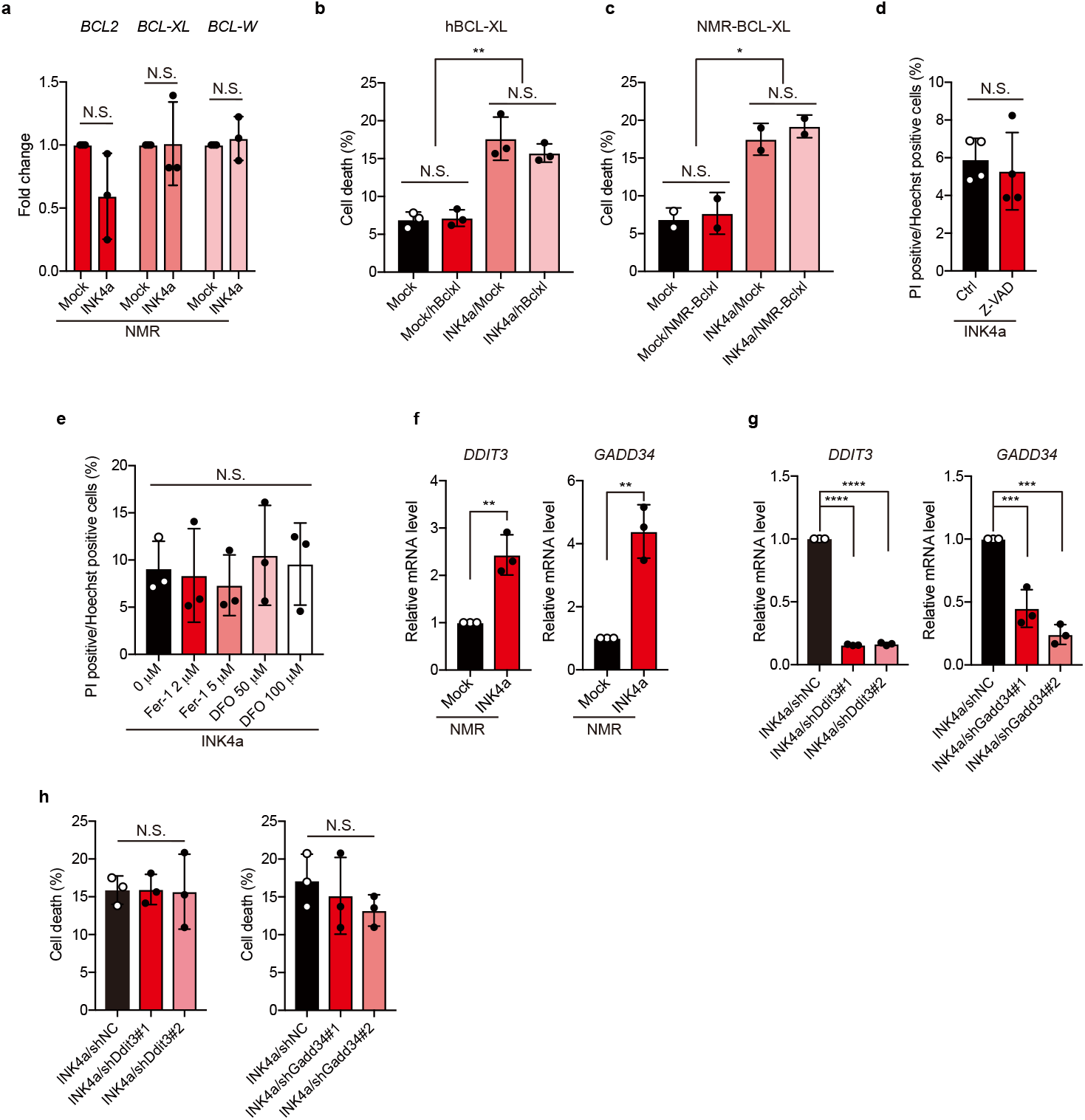
Overexpression of anti-apoptotic genes, treatment of ferroptosis inhibitors or inhibition of ER-stress do not suppress SCD. **a**, RNA-seq expression level analysis of anti-apoptotic BCL2 family genes. Y-axis: fold changes of BCL2 family mRNAs in INK4a-transduced NMR-fibroblasts relative to mock-transduced NMR-fibroblasts from n = 3 biological replicates. **b, c**, Quantification of Annexin V-positive cells in INK4a-tranduced NMR-fibroblasts with human-BCL-XL (*n* = 3 biological replicates) or NMR-BCL-XL (*n* = 2 biological replicates) transduction (%). **d**, 20 days after INK4a transfection into fibroblasts, the cells were treated with Pan-caspase inhibitor, Z-VAD-FMK for 24 h and PI-positive cells were quantified (%). *n* = 4 biological replicates. **e**, 20 days after INK4a transfection into fibroblasts, the cells were treated with indicated doses of Fer-1 or DFO for 2h and PI-positive cells were quantified (%). **f**, qRT-PCR analysis for the expression of *DDIT3* and *GADD34* in INK4a-transduced NMR-fibroblasts. **g**, qRT-PCR analysis for the expression of *DDIT3* and *GADD34* in INK4a-transduced NMR-fibroblasts transduced shRNA for each gene. **h**, Quantification of Annexin V-positive cells transduced shRNA for each gene. * *P* < 0.05; ** *P* < 0.01; *** *P* < 0.001; **** *P* < 0.0001; *t*-test for **a, d**, and **f**; one-way ANOVA followed by Tukey’s multiple comparison test for **b** and **c**; one-way ANOVA followed by Dunnett’s multiple comparison test for **e, g** and **h**. Data are mean ± SD from *n* = 3 biological replicates for **a, b, e, f, g** and **h**.

**Supplementaly Table 1:**
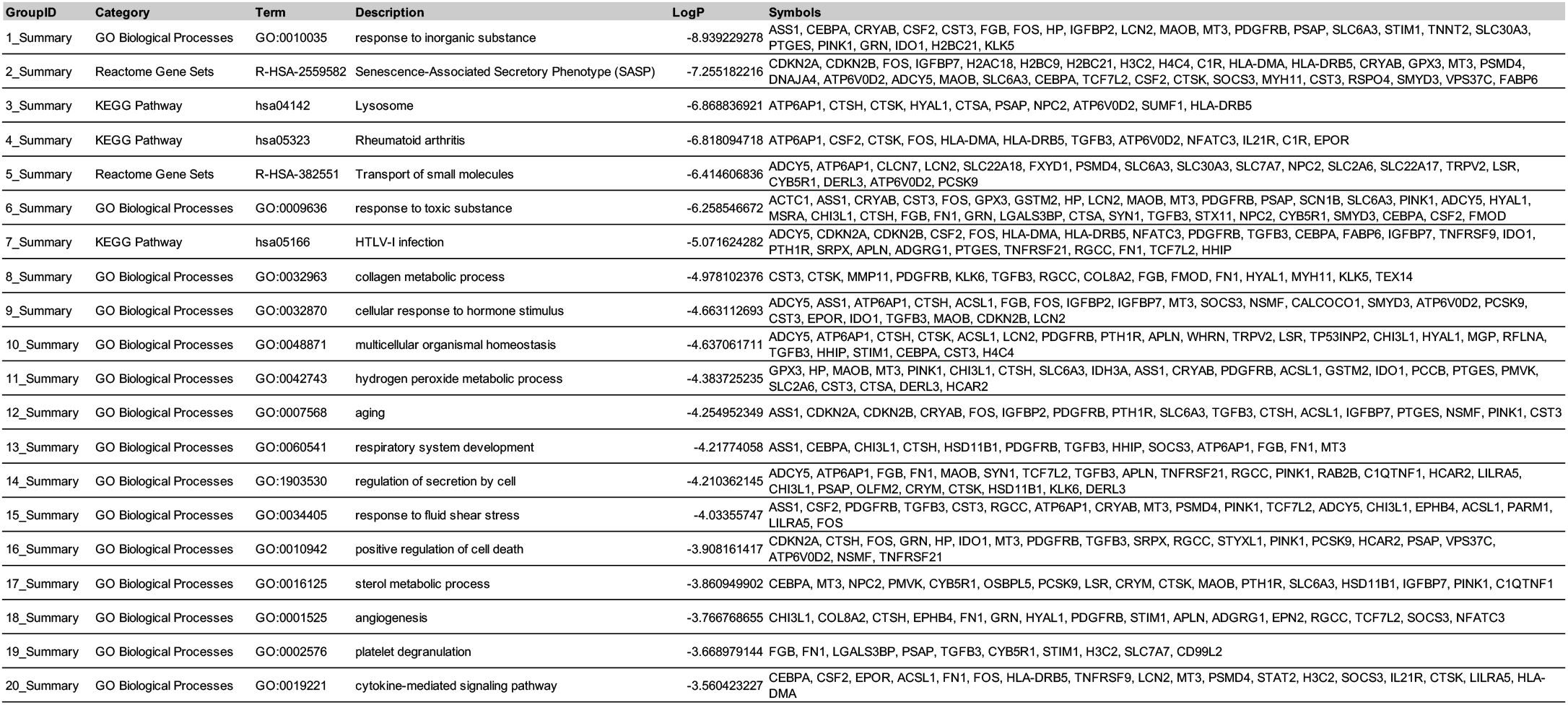
Summary of enriched GO terms and pathways.

**Supplementaly Table 2:**
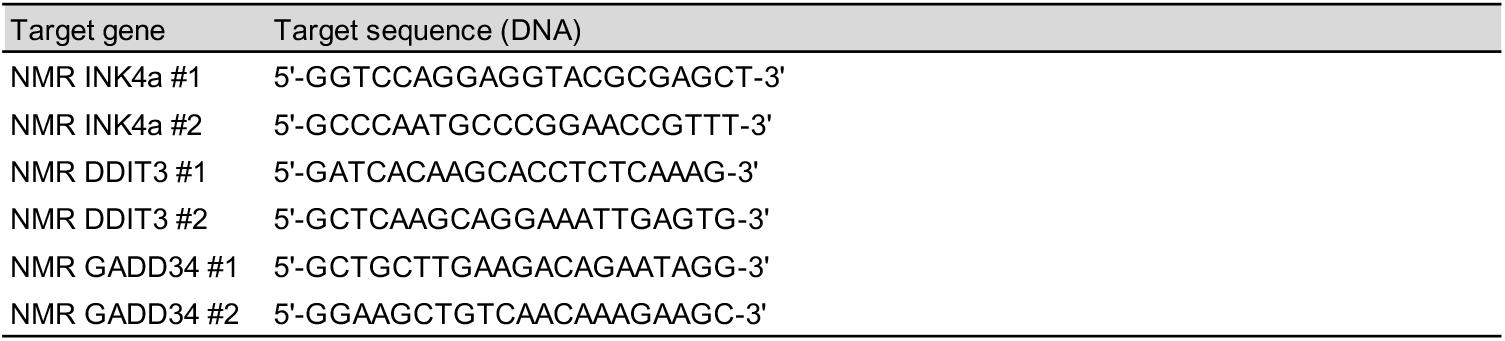
shRNA sequences.

**Supplementaly Table 3:**
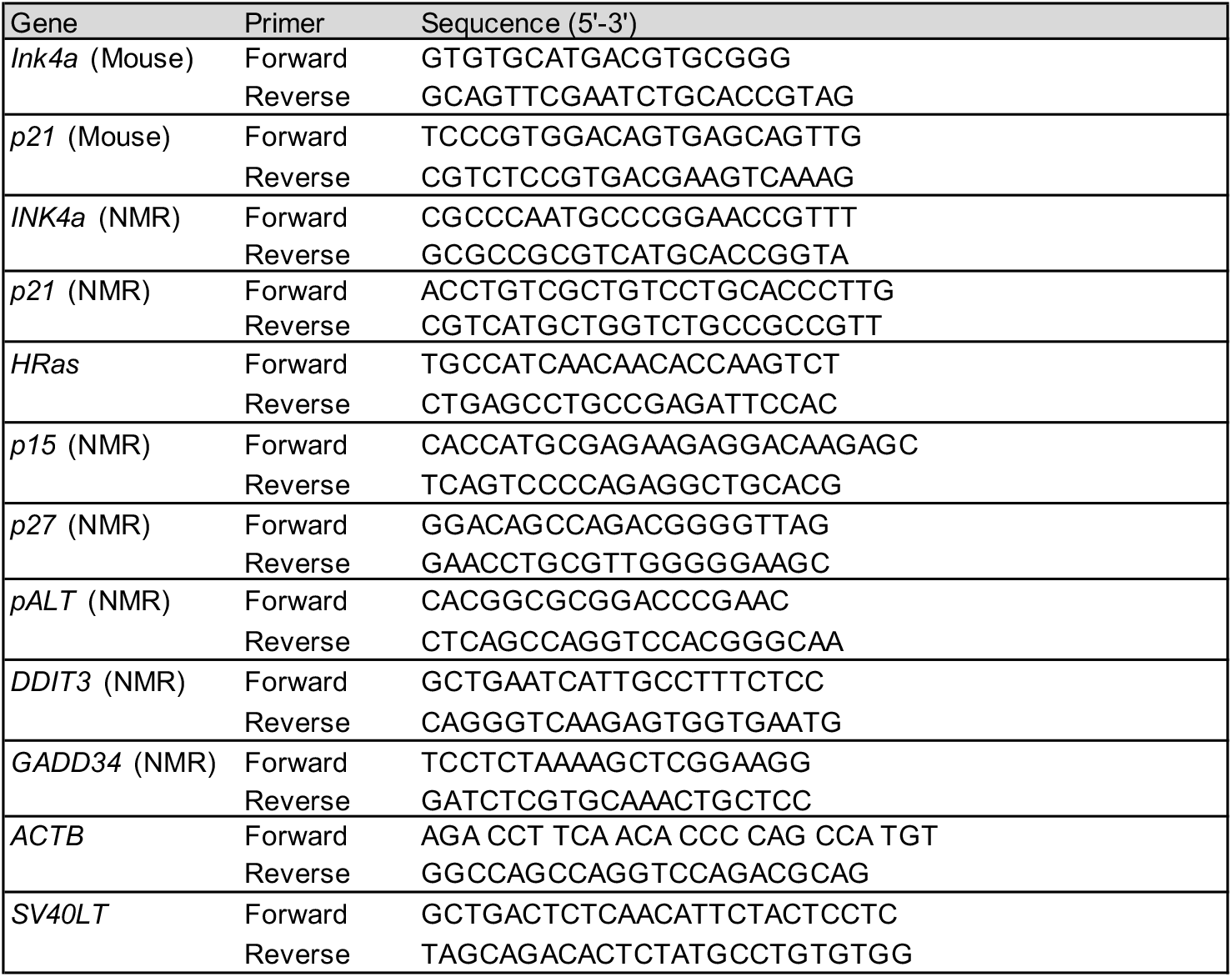
Primers used in this study.

